# CoCo: RNA-seq Read Assignment Correction for Nested Genes and Multimapped Reads

**DOI:** 10.1101/477869

**Authors:** Gabrielle Deschamps-Francoeur, Vincent Boivin, Sherif Abou Elela, Michelle S Scott

## Abstract

**Motivation:** Next generation sequencing techniques revolutionized the study of RNA expression by permitting whole transcriptome analysis. However, sequencing reads generated from nested and multi-copy genes are often either misassigned or discarded, which greatly reduces both quantification accuracy and gene coverage.

**Results:** Here we present CoCo, a read assignment pipeline that takes into account the multitude of overlapping and repetitive genes in the transcriptome of higher eukaryotes. CoCo uses a modified annotation file that highlights nested genes and proportionally distributes multimapped reads between repeated sequences. CoCo salvages over 15% of discarded aligned RNA-seq reads and significantly changes the abundance estimates for both coding and non-coding RNA as validated by PCR and bed-graph comparisons.

**Availability:** The CoCo software is an open source package written in Python and available from http://gitlabscottgroup.med.usherbrooke.ca/scott-group/coco.

## 1 Introduction

Detection and quantification of RNA transcripts is a critical step to understand the mechanism of gene expression and its impact on cell function. Traditionally, transcript abundance has been evaluated using techniques that target one known RNA sequence at a time, as in the case of quantitative RT-PCR. More recently, the development of RNA-sequencing techniques (RNA-seq) revolutionized transcriptome analysis by providing the tools necessary to study, at least in theory, all RNA transcripts simultaneously. Diverse library preparation protocols exist, the most commonly used ones focusing on particular classes of RNA through enrichment steps. Such strategies include polyA enrichment, non-rRNA enrichment (e.g. rRNA depletion), small RNA enrichment and enrichment for RNAs bound to specific factors (Conesa, et al., 2016; Hrdlickova, et al., 2017; O’Neil, et al., 2013). All these protocols detect certain levels of non-coding RNA. Strategies that employ rRNA depletion are the best approaches to detect a wide range of different classes of both coding and non-coding RNAs including lncRNAs (long non-coding RNA), small nuclear RNAs (snRNAs) and 7SL RNA (Boivin, et al., 2018; Lai, et al., 2016). However, no matter how the sequencing library is created, the capacity to correctly quantify RNA abundance ultimately depends on the correct assignment of the sequencing reads, including those generated by non-coding RNA.

Accurate RNA quantification is not easy to achieve since it depends on the quality of the transcriptome annotation used as a reference, the complexity of the target RNA sequence and its genomic context. RNA quantification is particularly difficult in the case of small and mid-size non-coding RNA, which are often produced from multiple genes and/or nested in other genes (Boivin, et al., 2018; Luo and Li, 2007; Mohammed, et al., 2014; Weber, 2006). According to Ensembl annotations (Yates, et al., 2016), in human, 2596 genes, including miRNA, scaRNA (small Cajal-body RNAs), snRNA, snoRNA (small nucleolar RNAs), tRNA (transfer RNAs) and lncRNA are located in introns and many overlap exons (**Supplementary Table 1**). In total, 1838 protein-coding genes, lncRNA and pseudogenes host or overlap smaller non-coding RNA (**Supplementary Table 2**). While some non-coding RNA are expressed from independent promoters like tRNA (Paule and White, 2000) many others do not have independent promoters and are at least partially linked to the expression of their host gene (Boivin, et al., 2018; Filipowicz and Pogacic, 2002; Matera, et al., 2007). Regardless of the mode of expression, correct identification of reads from nested and multi-copy genes is essential for the accurate quantification of both coding and non-coding RNA.

**Table 1.** Comparison of runtime and memory usage for available read assignment tools for a dataset of 41.5 M read pairs with a computer having 32 GB of available RAM and 24 cores.

| Tool | CPU time <sup>a</sup> | Walltime <sup>a</sup> | RAM (MB) | Threads |
| --- | --- | --- | --- | --- |
| CoCo | 1:40:08 | 0:49:26 | 6634 | 24 |
| rsem | 20:30:37 | 3:50:13 | 30372 | 24 |
| featureCounts<br>standard | 0:06:44 | 0:01:57 | 973 | 24 |
| featureCounts<br>optimized | 0:07:15 | 0:01:15 | 606 | 24 |
| HTSeq-count <sup>b</sup> | 4:51:14 | 3:10:46 | 173 | 24 & 1 |
| Cufflinks <sup>b</sup> | 15:17:47 | 14:02:29 | 25432 | 24 |

<sup>a</sup> CPU time and walltime are measured in hours:minutes:seconds.
<sup>b</sup> Cufflinks and HTSeq require sorting before the read assignment can occur. The values given include sorting before the tool is run.

Abundance estimation from RNA sequencing data generally requires a pipeline that aligns reads to a reference genome and then assigns them to annotated genes (Conesa, et al., 2016). In higher eukaryotes, mapping RNA reads to a genome requires a gapped alignment to avoid the DNA intronic sequences, which is accurately performed by standard software like the splice-aware aligners STAR (Dobin and Gingeras, 2016) and HISAT (Kim, et al., 2015). Once aligned, the sequencing reads must be assigned to genes, a task made challenging by overlapping genes and multimapped reads. Indeed, many available tools and commonly used settings only quantify correctly genes generating uniquely mapped reads, and reads mapping to more than one locus are typically discarded (**Fig. 1**). Reads originating from non-coding RNAs overlapping exons (mainly retained introns) of longer protein-coding RNA or lncRNA are either wrongly assigned to their host gene or labelled as ambiguous and discarded (**Fig. 1**). In addition, reads originating from duplicated genes are often discarded by default and many such genes, that we term multimapped genes, are under-represented in RNA-seq assigned counts.

**Figure 1.**
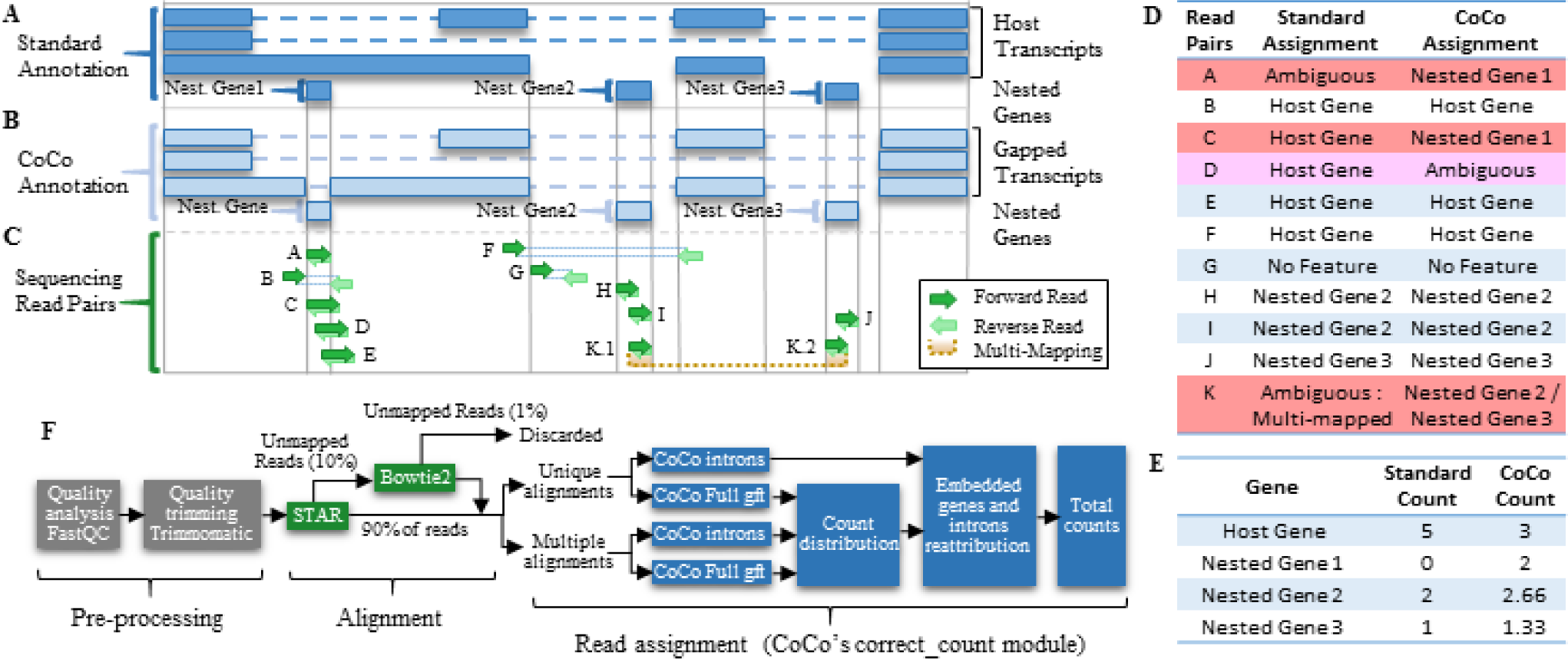
CoCo read correction scheme for nested and multimapped genes. (A) Representation of a standard gene annotation used for depicting a genetic locus containing one host gene and three nested genes. The dashed lines indicate introns while the dark blue boxes indicate exons. (B) Representation of the gene annotation produced using the correct_annotation module of CoCo showing a gap in the retained intron over the first nested gene. (C) Examples of potential read pairs overlapping the different features and multimapped read pairs. (D) Comparison of the read pair assignment using standard and CoCo pipelines, for each of the read pairs illustrated in C. The reads that are differentially assigned by CoCo are highlighted in red. (E) Comparison of the read count estimates by the standard and the CoCo pipelines, based on the assignments listed in D. (F) Flow chart of the CoCo pipeline. Pre-processing and alignment steps are shown before the correct_count module application. The correct_count module then assigns reads with Subread’s featureCounts using the gapped CoCo annotation (built with the correct_annotation module). Read pairs resulting in multiple alignments are considered separately and distributed proportionally to the uniquely assigned read pairs.

To increase the overall accuracy of RNA quantification and monitor the expression pattern of overlapping and repetitive genes, we developed a <u>co</u>unt <u>co</u>rrector (CoCo) pipeline that rescues and correctly assigns otherwise ambiguous sequencing reads. CoCo employs the read assignment function featureCounts from Subread (Liao, et al., 2013), providing it with a modified annotation file that highlights nested genes, while ensuring appropriate distribution of multimapped reads. CoCo salvages and/or reassigns over 15% of aligned RNA-seq reads, significantly changing the abundance estimates for several classes of RNA and thus providing insight into the expression dynamics of often ignored classes of repetitive and overlapping genes. We investigated the performance of the main available read assignment pipelines, comparing their accuracy to digital PCR abundance values and bedgraph estimates, showing that CoCo performs best using these metrics while requiring reasonable runtime and memory.

## 2 Methods

### Sequencing datasets

To ensure representative abundance of transcripts from nested and multimapped genes, most of which are highly structured RNAs, we chose TGIRT (thermostable group II intron reverse transcriptase) sequencing datasets. TGIRT displays high processivity and fidelity, providing an accurate picture of the ribodepleted transcriptome (Boivin, et al., 2018; Nottingham, et al., 2016; Qin, et al., 2016). The samples considered are GSM2631741/GSM2631742 (fragmented) as well as GSM2631743/ GSM2631744 (not fragmented) from GEO series GSE99065. While fragmented datasets enable the study of the whole transcriptome, non-fragmented datasets emulate size-selecting short RNAs (Boivin, et al., 2018).

### Read alignment

FastQC was used to check Fastq files for quality. Reads were trimmed using Cutadapt (Martin, 2011) (using parameters -g GATCGTCGGACTGTAGAACTCTGAACGTGTAGATCTCGGTGGT CGCCGTATCATT -a AGATCGGAAGAGCACACGTCTGAACTCCAGTCACATCACGATCTCGTATGCCGTCTTCTGCTTG --mini-mum-length 2) and Trimmomatic (Bolger, et al., 2014) (with TRAILING:30) to remove adaptors and portions of reads of low quality, respectively. Read pairs were then aligned to the human genome build hg38 using an annotation file obtained from Ensembl (described below) with the splice-aware RNA-seq aligner STAR (Dobin and Gingeras, 2016) using the following parameters: --outSAMtype BAM SortedByCoordinate, --outSAMprimaryFlag AllBestScore, --alignIntronMax 1250000, all other parameters at default values. Reads not aligned using STAR were aligned once again using Bowtie v2 (Langmead and Salzberg, 2012), which performs well for the alignment of shorter reads. Parameters used for Bowtie were the following: --local, -p 24, -q, -I 13. Read pairs successfully aligned by STAR or Bowtie were merged into a BAM file and separated into two groups: those that align to one genomic position and those that align to more than one genomic position (**Fig. 1F**).

### Read assignment and corrections

The correct_count module from CoCo uses featureCounts (Liao, et al., 2013) to assign aligned read pairs to their corresponding genes by their genomic coordinates using the given annotation file as reference (**Fig. 1**). The following featureCounts parameters are used by CoCo: (-C, --largestOverlap, -p, -B, -s 1, --minOverlap 10, [-M]). The parameter -M is not specified when assigning uniquely mapped reads and is specified when assigning multimapped reads. CoCo uses a corrected annotation produced from Ensembl supplemented annotation (described below) by removing regions corresponding to nested genes from longer host genes (described below).

Read pairs aligning to only one genomic location increase the alignment count by one for the genomic feature encoded at that location. By default, correct_count distributes the reads between their assigned genes according to their number of assigned uniquely mapped reads. If no singly mapped reads exist for any of the assigned genes, the reads are distributed uniformly. Total counts assigned to a gene correspond to the sum of the uniquely aligned read pairs and the proportion of multimapped read pairs assigned to the gene. The correct_count module also corrects for over-attribution of reads to nested genes by subtracting the length normalized background counts attributed to the feature in which the nested gene is encoded, as described in **Supplementary Fig. 1**.

### Annotation supplementation

An annotation file in gene transfer format (.gtf) was obtained from Ensembl (Yates, et al., 2016) (hg38, v87). The annotation file was supplemented with 628 additional tRNA from GtRNAdb (Chan and Lowe, 2016) and with 20 snoRNA from Refseq (O’Leary, et al., 2016) that were missing from Ensembl annotations. In addition, 63 gene annotations were removed from the gtf file because they overlap another gene over more than 90% of their length and keeping them would result in reads aligning to them being labelled as ambiguous. These 63 redundant genes consist of 17 snoRNAs, 43 miRNAs, 2 lincRNAs and 1 antisense RNA. Details of added and removed genes are given in **Supplementary Data File 1**.

### CoCo’s annotation correction

The correct_annotation module of CoCo produces a modified annotation file in which any exon position overlapping a snoRNA, scaRNA, snRNA, tRNA or miRNA is removed, resulting in a gapped annotation file (**Fig. 1B**). To do so, correct_annotation builds a list of gene coordinates corresponding to the above biotypes and then seeks overlaps with exons from all transcripts of genes from other biotypes using Bedtools intersect (Quinlan and Hall, 2010). Exons overlapping a nested gene have their overlapping positions removed, effectively splitting the exon in two if the nested gene is fully within the exon, or truncating the exon if the nested gene overlaps its end. From this, two new.gtf annotation files are made, one containing all the genes and transcripts, the other one including only the portion of the genes hosting a nested gene (respectively referred as “CoCo Full gtf” and “Introns gtf” in **Fig. 1** and **Supplementary Fig. 1**)

### Conversion from counts to TPM

The counts obtained from CoCo’s correct_count module were normalized by the length of the main transcript of the gene and the read count to give transcripts per million (TPM) as described further in (Boivin, et al., 2018):

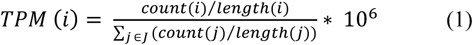

where count(i) and length(i) represent respectively the number of read pairs aligning to gene i and the length of gene i, and J represents the set of all genes in the annotation. The final output of the correct_count module holds the raw read counts, the CPM and the TPM values associated to annotated genes. **Supplementary Data File 2** gives average abundance values in TPM, estimated using the CoCo pipeline as well as all other read assignment tools considered, for all genes.

### Alignment Visualization

BAM files were converted to Bedgraph files using CoCo’s correct_bedgraph module (**Supplementary Fig. 2**) which uses pairedBamToBed12 (https://github.com/Population-Transcriptomics/pairedBamToBed12) and bedtools genomecov (v2.25.0) to produce a bedgraph of the distribution of the reads of a paired-end dataset. The bedgraphs were visualized using the Integrative Genomics Viewer IGV (Robinson, et al., 2011) with the hg38 human genome build and the Ensembl annotation tracks.

### Gene biotype pooling

Gene biotype groups “Protein_coding”, “Pseudogene” and “Long_noncoding” were pooled as recommended by Ensembl (http://ensembl.org/Help/Faq?id=468). The group “Other” corresponds to all other biotypes not listed.

### Comparison to other read assignment tools

CoCo was compared to the following read assignment tools: featureCounts (Liao, et al., 2013), HTseq-count (Anders, et al., 2015), RSEM (Li, et al., 2010), Cufflinks (Trapnell, et al., 2012) and STAR (Dobin and Gingeras, 2016). The parameter values used for each tool are indicated in **Supplementary Table 4**. Throughout the text, featureCounts with typical parameter values is used as the standard read assignment pipeline.

### PCR

Digital PCR was performed by the Université de Sherbrooke Rnomics Platform (http://rnomics.med.usherbrooke.ca/). Droplet Digital PCR (ddPCR) reactions were prepared using 10ul of 2X QX200 ddPCR EvaGreen Supermix (Bio-Rad), 10 ng (3 µl) cDNA,100 nM final (2 µl) primer pair solutions and 5ul molecular grade sterile water (Wisent) for a 20ul total reaction. Each reaction mix was converted to droplets with the QX200 droplet generator (Bio-Rad). Droplet-partitioned samples were then transferred to a 96-well plate, sealed and cycled in a C1000 deep well Thermocycler (Bio-Rad) under the following cycling protocol: 95°C for 5 min (DNA polymerase activation), followed by 50 cycles of 95°C for 30 s (denaturation), 59°C for 1 min (annealing) and 72°C for 30 s (extension) followed by post-cycling steps of 4°C for 5 min and 90°C for 5 min (Signal stabilization) and an infinite 12°C hold. The cycled plate was then read using the QX200 reader (Bio-Rad). The concentration reported is in copies/ul of the final 1x ddPCR reaction (using QuantaSoft software from Bio-Rad). All primer sequences are available in **Supplementary Table 3**.

## 3. Results

### The CoCo correction for nested genes and multimapped reads

Standard gene quantification programs assign reads according to the amount of overlap between the read and the feature being quantified. As a consequence, reads mapping with the same number of matches to a host gene and to a small non-coding RNA gene nested within its intron/exon are often considered ambiguous (e.g. **Fig. 1 C,D**, read pair A). In addition, reads from nested genes that exceed the annotations (which, in many cases, are not accurate (Deschamps-Francoeur, et al., 2014; Kishore, et al., 2013)), by even only one nucleotide, are typically automatically assigned to the host gene (e.g. **Fig. 1C,D**, read pair C). To address these problems, we have developed the Count Corrector (CoCo) package, which consists of three main modules: 1) the correct_annotation module which generates gapped annotation files in which the regions of the host gene transcript features overlapping with nested genes are precisely removed (**Fig. 1B**), 2) the correct_count module which recuperates the reads associated with nested and multimapped genes using the modified annotation (**Fig. 1D** and **E**), and 3) the correct_bedgraph annotation which produces accurate representations of paired-end reads (**Supplementary Fig. 2**).

To test the quantification accuracy of the CoCo pipeline, we examined its capacity to correctly assign and quantify sequencing reads using four RNA-seq datasets, and compared its quantification to those of the main read assignment pipelines available. All four samples were generated using a library preparation protocol that accurately detects both coding and non-coding RNA classes, thanks to the use of the TGIRT reverse transcriptase and ribodepletion (Boivin, et al., 2018). Two of the samples were fragmented which enables the quantification of both long and short RNAs (for example nested genes and their host gene) while the other two samples were not fragmented, providing a deeper view of short RNAs, which is comparable to size selection RNA-seq. Following standard pre-processing steps, reads were mapped to the hg38 human genome using STAR, resulting in an average alignment rate of 90%. The 10% unaligned reads using STAR are mostly alignments deemed ‘too short’ by STAR and thus a second alignment step using Bowtie was employed, resulting in an overall average alignment rate of 99% (**Fig. 1F**). Aligned reads were then assigned to annotated genes using the correct_count module of the CoCo pipeline and the annotation files provided by CoCo’s correct_annotation module. The correct_count module not only reattributes reads from host genes to nested genes but also corrects the reassignment considering the background read counts of the feature in which the gene is embedded (**Supplementary Fig. 1**), as discussed below.

### Impact of CoCo on the quantification of nested genes and comparison to other read assignment tools

To evaluate the impact of CoCo on RNA detection and quantification, we compared the aligned read assignments obtained using the CoCo pipeline to values obtained using five currently available and commonly used read assignment pipelines, described in the Methods and in **Supplementary Table 4**. The pre-processing and alignment steps (described in the Methods and in **Fig. 1F**) were the same for all read assignment tools, except for RSEM which requires an alignment to the transcriptome instead of the genome (so STAR but not bowtie was used, with the additional parameter --quantMode TranscriptomeSAM). As indicated in **Fig. 2** and **Supplementary Figs. 3-7**, the CoCo pipeline performs well in the quantification of nested genes, both in fragmented and non-fragmented datasets, always obtaining raw read counts in close agreement with those estimated from the bedgraph, for all nested genes considered. In contrast, HTSeq-count, featureCounts with standard parameter values and STAR fail to detect many nested genes, in all datasets considered. Cufflinks was not considered for this comparison because it does not provide raw read counts that can be compared to bedgraph counts. Nested genes that are poorly or more often completely non-detected by HTSeq-count, featureCounts with standard parameter values and STAR include SNORD55, SNORD38B (**Fig. 2**), SNORD14D, SNORD14C (**Supplementary Fig. 3**), RNU5E-6P (**Supplementary Fig. 4**), SNORA58 (**Supplementary Fig. 5**), tRNA-Gln-TTG-2-1 (**Supplementary Fig. 6**) and SCARNA15 (**Supplementary Fig. 7**), which all overlap, partially or completely, retained introns or exons of their host gene. Manual inspection of the alignment file and visual inspection of the bedgraphs confirm that all these nested genes have corresponding aligned reads and should thus be detected and quantified as expressed. RSEM and featureCounts with optimized parameter values perform better than HTSeq-count, STAR and featureCounts with standard parameter values, but still deviate more from bedgraph estimates than does CoCo (see for example **Supplementary Figs. 3, 4, 6 and 7**). In the presence of ambiguous reads (**Fig. 1, read pair A**), featureCounts_optimized splits the read counts equally between the nested gene and the host gene. For example, in the case of the PTCH2 gene locus (**Supplementary Fig. 4**), a visual inspection of the bedgraph indicates that the great majority if not all read counts should be attributed to the nested gene RNU5E-6P while featureCounts_optimized splits the read counts equally between the two genes for the fragmented dataset. Interestingly, in this case, CoCo not only corrects the abundance values of the nested gene but also adjusts the counts given to the host gene, reassigning 89% of the counts (313 read pairs out of 350) from the host gene to the snRNA. Based on these results, we conclude that CoCo is capable of rescuing and properly assigning reads from the nested genes lost to several read assignment tools and generates quantification values that are most consistent with the bedgraph profiles.

**Figure 2.**
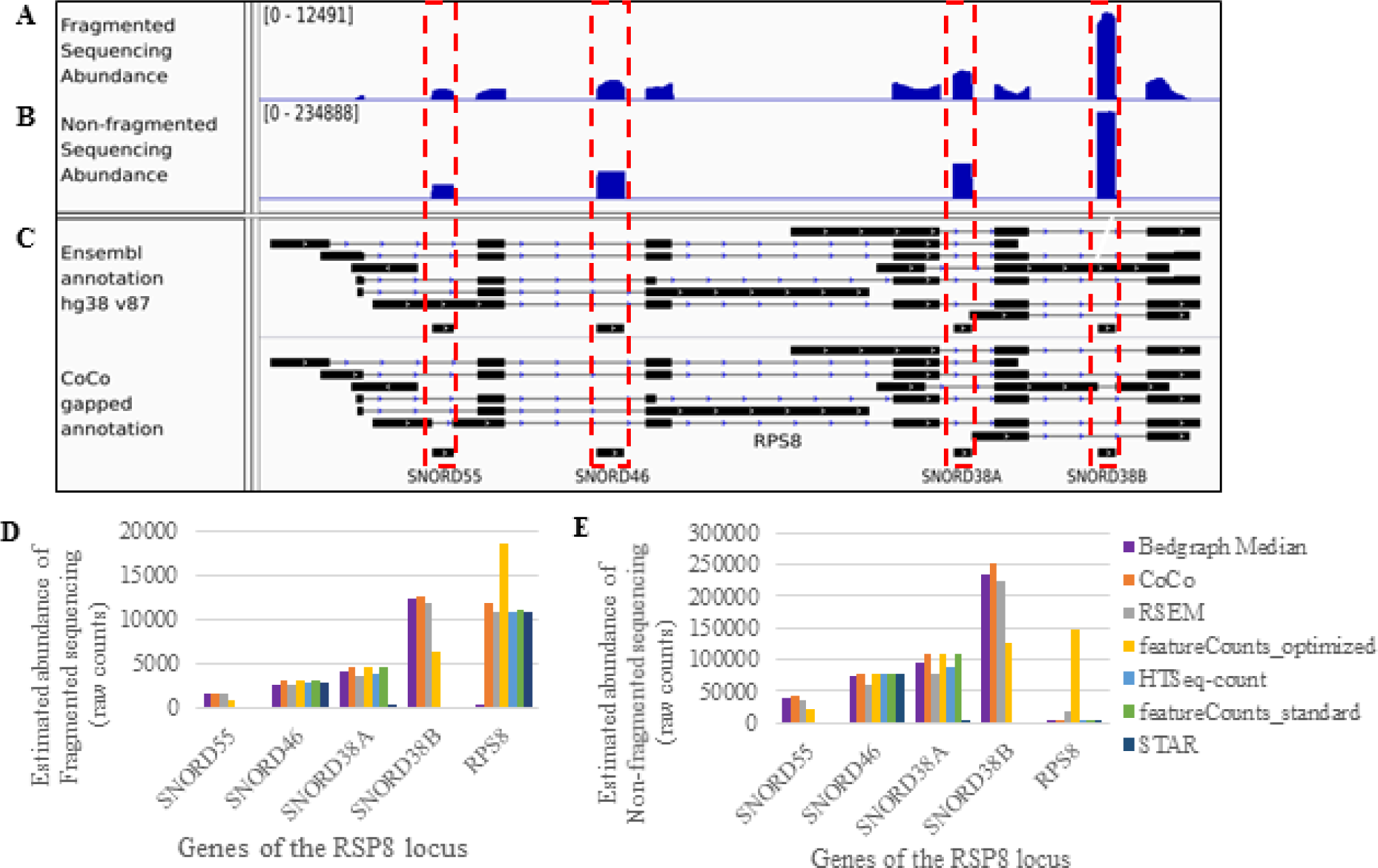
Examples of bedgraphs illustrating the CoCo quantification correction for nested genes. Example of a host gene holding multiple intron-encoded snoRNAs. The ribosomal protein gene RPS8 harbours four box C/D snoRNAs in four separate introns, two of which are retained in certain RPS8 splice variants. A screenshot of the sequencing abundance tracks is shown for fragmented (A) and non-fragmented (B) datasets, below which is shown the annotation tracks including the original Ensembl annotations and the CoCo gapped annotation (C). Exons are represented as boxes, introns as lines with arrows and nested small RNA genes are highlighted with red dashed boxes. (D,E) Histogram showing the read pair raw counts given by CoCo and other read assignation tools for the snoRNA and host genes illustrated in A-C, for the fragmented (D) and non-fragmented (E) datasets.

### Background correction for nested genes

In order to accurately quantify read counts from nested genes and distinguish their reads from the host gene reads, CoCo carries out a background correction. This is achieved by subtracting the average read count of the feature in which the nested gene is encoded from the read count assigned to the nested gene. This correction does not significantly change the final read counts of genes nested in host gene features (most often introns) that are expressed at low levels, (**Supplementary Fig. 8)**. However, this correction has a major impact when the host gene feature encoding the nested gene is highly expressed. For example, in the case of the CH507-513H4.1 locus which hosts miRNAs miR-3648 and miR-3687, the reads were originally attributed to the miRNA despite the absence of corresponding peaks in the bedgraph. This inappropriate assignment is no longer observed following background correction (**Supplementary Fig. 9**).

### Quantification of transcripts from duplicated genes

A large number of genes, in particular those producing non-coding RNA, exist in more than one copy. The RNA produced from most of these different forms of repeated features cannot be quantified by standard read assignment modules due to their identical or near identical sequences that result in multimapping of sequencing reads (e.g. **Fig. 1C,** read pair K). The CoCo pipeline recuperates these otherwise lost reads by distributing the counts between all genes assigned according to the distribution of their uniquely mapped read pairs when possible. For small non-coding RNAs such as snoRNAs, uniquely mapped reads typically originate from flanking genomic sequences that are seldom included in the transcripts. Longer genes are more likely to have longer proportions of unique sequences. For example, as indicated in **Supplementary Fig. 10**, the annotated mature forms of SNORD103A and SNORD103B are 100% identical. As a result, 96% (2314/2401) of all read pairs aligning to these snoRNAs map equally well to both SNORD103A and SNORD103B and would thus be discarded without considering multimapped reads. Considering the distribution of the 87 uniquely aligned read pairs (to non-identical genomic flanking sequences for SNORD103A and SNORD103B), CoCo distributed the 2314 multimapped read pairs proportionally to the uniquely mapped read pairs (**Supplementary Fig. 10B**). As a consequence, the sum of the read counts to the SNORD103 family increased by 25X passing from 87 (considering only uniquely mapped reads) to 2401 read pairs (using the CoCo’s correct_count module). Therefore, by using the uniquely mapped reads as a guide, CoCo rescues and re-distributes the multimapped reads to provide a realistic read distribution. Overall, the CoCo multimapped correction module increased the estimated abundance of 1443 multimapped genes, most of which are tRNAs and snoRNAs, by more than two-fold.

### Experimental validation of CoCo based quantification for nested and repeated genes

To experimentally validate the accuracy of the CoCo based quantification, we chose nine overlapping or repeated genes and examined their abundance using droplet digital PCR (ddPCR), comparing these values to sequencing abundance estimates. These 9 genes include 7 snoRNAs overlapping with protein coding genes and 2 repeated gene families (RN7SK and RN7SL2). In the case of the multimapped genes RN7SK and RN7SL2, their abundance was estimated to be respectively 2 and 4 times higher respectively (**Fig. 3**) using the CoCo pipeline compared to feature-Counts_standard estimates. When the ddPCR was compared to all read assignment tools considered (**Fig. 3** and **Supplementary Fig. 11**), as for the bedgraph comparison, HTSeq, STAR and featureCounts using standard parameter values did not detect the nested genes while CoCo, RSEM and featureCounts with optimized parameter values agreed with ddPCR values, obtaining Pearson correlation values above 0.99.

**Figure 3.**
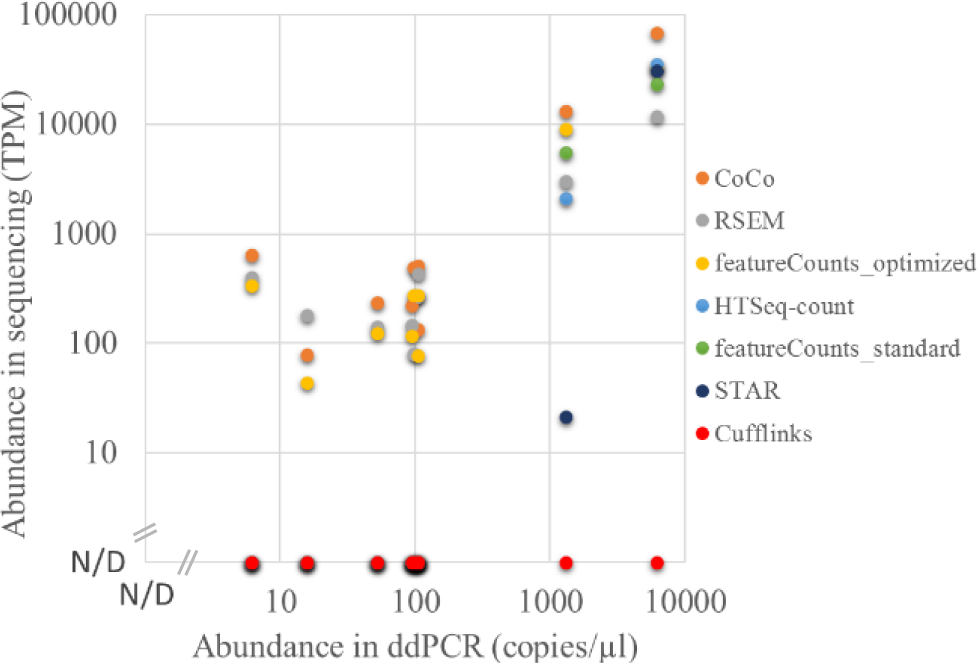
Comparison of read assignment tools with ddPCR abundance estimates. Scatter plots showing abundance values obtained by ddPCR compared with TPM estimates of the fragmented datasets from read assignment tools considered. N/D: “Not Detected”.

### CoCo increases the number of sequencing reads considered and enhances the detection of non-coding RNA

The capacity to detect transcripts from repeated and overlapping genes not only provides new information about the expression dynamics of these genes but may also alter the overall distribution of read counts. Accordingly, we examined the impact of using CoCo on the overall distribution of transcript estimates in the human genome. Comparison of the read assignments obtained using the CoCo pipeline or the standard featureCounts pipeline indicates that while 65.8% of the read pairs in fragmented datasets were assigned identically by both strategies and 18.6% were assigned by neither, more than 15% were only assigned by CoCo (**Supplementary Fig. 12**). Most of the read pairs correctly assigned by the standard pipeline originate from protein-coding genes (74%) and tRNAs (12%) (**Supplementary Fig. 12E left panel**). Read pairs that were not assigned by either pipeline originate mostly (88%) from currently unannotated genomics regions (**Supplementary Fig. 13**), as is the case for example for an intronic region in the AKAP6 gene in which >32000 read pairs align but no feature is annotated (**Supplementary Fig. 14**).

Most of the read pairs uniquely assigned by CoCo (15.7%) originate from multimapped genes, representing 12.9% of all aligned read pairs (**Supplementary Fig. 12A**). As expected, the rescued multimapped read pairs originate mainly from non-coding RNA including tRNAs (37%), 7SL (27%), snRNAs (14%) (**Supplementary Fig. 12C**). However, 13% of the reads aligned to protein-coding genes suggesting that there is a substantial number of protein-coding genes that contain repeated sequence (**Supplementary Fig. 12C**). A small proportion of the read pairs uniquely assigned by CoCo (2.4%) were aligned to overlapping genes that are labelled as ambiguous by standard pipelines (**Supplementary Fig. 12A** right panel). Most of the overlapping gene counts originated from non-coding RNA including snoRNAs (40%), snRNAs (37%) and 7SK (18%) (**Supplementary Fig. 12D**). In addition to the 15.3% of rescued ambiguous and multimapped read pairs, CoCo also reassigned a small proportion of reads (0.37%) that are misassigned by the standard pipeline (**Fig. 1C-D** read pair C and **Supplementary Fig. 15**). In all cases, misassignments by the standard pipeline result in erroneous association of reads to the host gene instead of the nested gene. SnoRNAs represent 94% of such reassignments (**Supplementary Fig. 15**). Together, these observations indicate that CoCo greatly increases the percentage of usable read counts and reduces the number of misassignments.

### CoCo provides a more accurate depiction of the transcriptome landscape

As a consequence of the CoCo correction, the overall distribution of the human transcriptome was modified considerably. The proportion of all read counts attributed to protein-coding genes in fragmented datasets was reduced by 12%, representing 74% of all assigned reads using the standard pipeline but only 62% using CoCo (compare **Supplementary Fig. 12E** left and right panels). In contrast, non-coding RNA including tRNAs and snRNA gain 4% and 2% of the total read counts, respectively. The most dramatic changes in total read count distribution was observed with the highly redundant (multimapped) gene coding for the signal recognition particle RNA 7SL, which was increased from 2% using the standard pipeline to 6% using CoCo. Analyses of transcript abundance distribution, which take into consideration the transcript length, indicate that the biggest impact of CoCo is in adjusting the proportion of coding to non-coding RNA and increasing representation of RNA families with a strong prevalence of repeated genes. As indicated in **Supplementary Fig. 12F**, CoCo estimates of transcript abundance reduced the proportion of protein coding genes by 5% while tripling the proportion of the 7SL and increasing the proportion of snoRNA and snRNA by 2% each, when compared to the standard pipeline.

The impact of CoCo on transcript quantification is most visible in cases where RNA abundance changes from completely undetected using standard pipelines to abundantly detected with the CoCo pipeline. (e.g. **Fig. 2, 3, Supplementary Figs. 3,4,6,7**). The de novo detection of these genes gives a new view of an otherwise uncharted portion of the human transcriptome. Accordingly, we counted the number and biotype distribution of genes only detected using CoCo as compared to a standard read assignment pipeline to better understand their origin and contribution to the human transcriptome, using a fragmented dataset to consider the full RNA landscape. As shown in **Fig. 4A**, the most affected RNA family is the snoRNAs with 22% of genes (137 out of 628 expressed snoRNAs) detected only with the CoCo pipeline. The distribution of the corrected abundance of these 137 ‘invisible’ C/D and H/ACA box snoRNA genes is displayed in **Supplementary Fig. 16**. More than 50% of these snoRNAs (80/137) have a corrected abundance above 100 TPM, and are thus amongst the most abundant RNAs in the cell (indeed, only 1.8% of all expressed genes have cellular transcripts with abundance >100 TPM). Ten such snoRNAs are even detected with an abundance as high as >1000 TPM (e.g. SNORD26, SNORD14C). These data clearly indicate that failure to detect snoRNAs using standard assignment methods is not restricted to rare or lowly expressed RNA. On the contrary, it is the highly expressed genes that are often missed, most likely due to the fact that highly expressed non-coding RNA tend to originate from more than one gene or to be embedded in highly transcribed genes. In addition to snoRNAs, a smaller but still significant proportion of each of the other main classes of non-coding RNA is not detected using standard techniques including >10% of scaRNA and as many as 27 snRNA genes, 27 tRNA genes and 224 lncRNAs genes (**Fig. 4A**).

**Figure 4:**
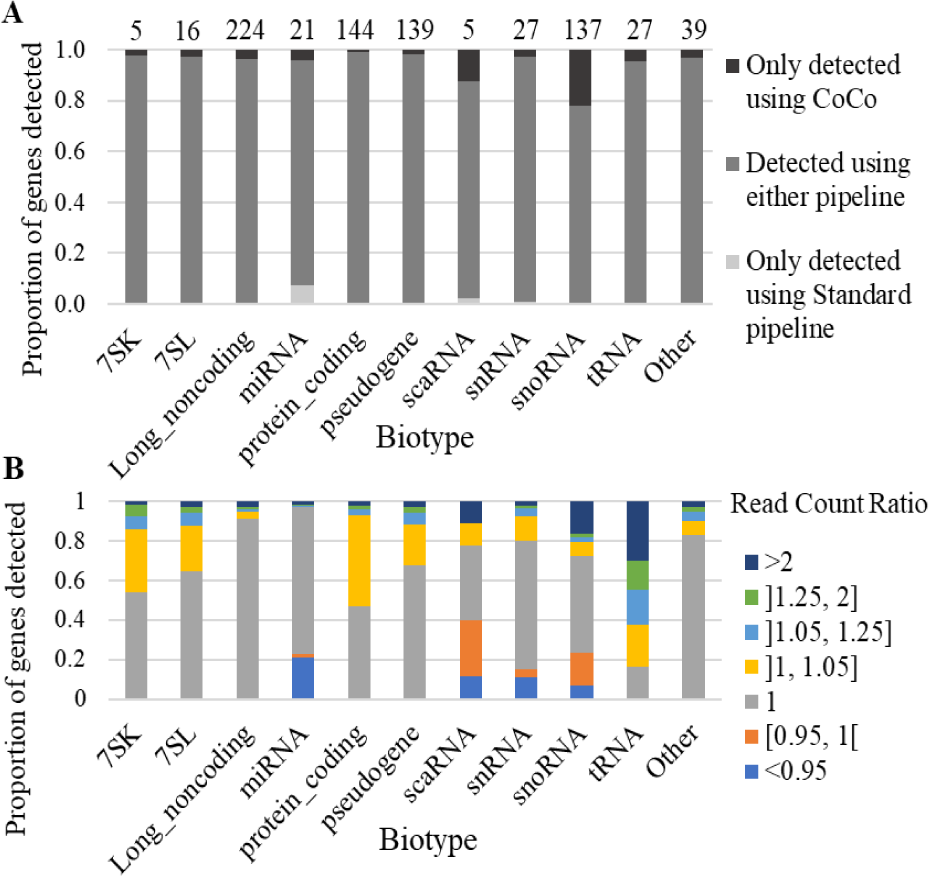
Effect of the CoCo pipeline on gene quantification of fragmented datasets by biotype. (A) The proportion of transcripts detected by either only the standard pipeline (light grey), both the standard and CoCo (intermediate grey) or only by the CoCo (dark grey) pipelines is shown as a bar graph. The number of genes only detected using CoCo is indicated at the top of the graph for each biotype considered. (B) Impact of CoCo on the read counts of different biotypes. Shown is a bar graph indicating the proportion of genes of each biotype displaying the indicated change in abundance following the CoCo correction.

A less obvious but equally important difference between the CoCo and standard pipelines lay in the accuracy of read assignments, especially to duplicated genes. For example, while both CoCo and the standard pipeline agree on tRNA being the most abundant transcript biotype (**Supplementary Fig. 12F**), this class of RNA features the largest number of genes with count corrections by CoCo. Indeed, 30% of tRNA genes more than doubled their abundance after using CoCo (**Fig. 4B**). Similarly, a great number of protein-coding genes (>50%) also increased in abundance, but unlike tRNA, the difference in read pair count was mostly modest with an average of 5% change in read counts (**Fig. 4B**). We therefore conclude that the correct counting and assignment of sequencing reads plays an important role not only in increasing the number of detected genes but in establishing the relative abundance of RNA within the transcriptome.

### Runtime and memory requirements

CoCo can be run using multiple threads thus improving its running performance. The runtime and memory required to use the different read assignment tools considered in this study is shown in Table 1.

## 4. Discussion

Increasing sequencing depth and coverage is considered the obvious target for improving the quality of *transcriptome* analysis. However, in this study, we show that the precision, quality and depth of transcriptome analysis can be greatly enhanced by carefully choosing and tuning read assignment tools using existing sequencing data. Modification of the read assignment pipeline for the analysis of a TGIRT-seq fragmented dataset, which provides an accurate view of the whole transcriptome (Boivin, et al., 2018) increased the proportion of assigned reads by 15% and modified the number of assigned reads for 50% of all expressed genes (**Fig. 4, Supplementary Fig.12**). Indeed, more than 750 additional transcripts were detected simply by correcting read association for nested genes (**Fig. 4A**), while multimapping affects the quantification of 15,121 genes. The problem with overlapping genes was solved by introducing gaps corresponding to nested gene positions in the reference annotation files (**Fig. 1,2 and Supplementary Figs.3-7**). The gaps ensure that read pairs are not automatically assigned to the host gene even if they slightly exceed the often inaccurate annotation of nested genes (Deschamps-Francoeur, et al., 2014; Kishore, et al., 2013). CoCo is not the only tool that can correctly address the quantification of nested genes, although CoCo provides abundance values that are most consistent with bedgraph estimates (examples shown in **Fig. 2** and **Supplementary Figs.3-7**) and performs well when compared to ddPCR (**Fig. 3**). Multimapped reads were dealt with by distributing them proportionally to uniquely mapped reads, as first introduced by MuMRescue (Faulkner, et al., 2008) and ERANGE (Mortazavi, et al., 2008). The corrections proposed in our study can be applied as a supplement to any read-assignment tool as a feature. By applying the correction tool CoCo, most sequence analysis pipelines will benefit from increased transcriptome coverage and more accurate transcript quantification.

Five different tools (one of which was run using two different settings) were chosen for comparison to the CoCo pipeline because they are widely used and easily implemented standalone pipelines. HTSeq-count, STAR, Cufflinks and featureCounts using standard parameter values performed poorly to quantify nested and multimapped genes, both according to comparisons to bedgraphs and comparison to ddPCR quantification. These four tools failed to detect many nested genes and did not accurately quantify many multimapped genes. In contrast, CoCo, RSEM and featureCounts with optimized parameter values performed much better at quantifying nested and multimapped genes, both according to bedgraph comparisons and ddPCR comparisons. RSEM’s greatest drawback is its long runtime and high memory usage while featureCounts with optimized parameter values does not assign ambiguous reads in an ideal manner as it simply splits the assignment counts equally for all features overlapping these reads. In addition, RSEM and featureCounts with optimized parameter values do not perform well when quantifying non-fragmented datasets, generally attributing too many reads to host genes. They are thus not recommended for size-selection datasets often used to quantify miRNAs. In general, if any of these tools are chosen to quantify RNA-seq datasets, we highly recommend the parameter values used in this study (see **Supplementary Table 4**) as using different parameter values can significantly change the quantification and thus the accuracy (compare for example the results obtained using featureCounts_standard and feature-Counts_optimized).

Accurately quantifying multimapped reads has been investigated in several studies. featureCounts annotates such reads, making it easy for CoCo’s correct_count module to distribute them according to uniquely mapped reads. featureCounts itself provides an option to deal with multimapped reads, by splitting them equally between all members of a multimapped group (using the –M, --fraction options). However, a significant proportion of these members are likely not expressed, particularly in the large families of ncRNAs with tens or even hundreds of copies, making CoCo’s strategy (the proportional distribution of multimapped reads) more realistic. Evaluating the accuracy of multimapped read assignment is difficult since unlike mRNA, most non-coding RNA lack external unique sequences that could be used to differentiate between genes with shared sequences. In building our pipeline for the quantification of nested and multimapped genes, we opted for a computationally quick and simple solution that enables the evaluation of the overall abundance of RNA generated from genes with shared sequence. While we cannot guarantee the accuracy of the abundance for each individual gene from a repeated family, the overall abundance of the RNA generated from these repeated families is accurately determined by CoCo as validated experimentally by ddPCR analysis and gene specific analyses (**Fig. 3** and (Boivin, et al., 2018)).

One of the most surprising observations in the application of CoCo is how many genomic regions do not have proper annotation and how this mis- or lacking annotation may affect the overall interpretation of transcript distribution. Indeed, while CoCo was able to recuperate the ambiguous reads due to gene overlap with small non-coding RNAs and multimapping, about 18.6% of all read pairs remained unassigned. Examination of these read pairs shows that the great majority align to unannotated regions in the genome (**Supplementary Fig. 14**), indicating that improving the annotations is now becoming essential to increase the coverage and the accuracy of RNA-seq quantification.

The read assignment corrections shown here are essential steps for studying the human transcriptome. Most of the multimapped and overlapping reads are generated from highly expressed genes and as such any changes in the read assignment of these genes will significantly affect the overall distribution of the human transcriptome. In addition, accurate quantification of nested and multimapped genes does not only enhance the detection of these types of RNA but also corrects the quantification of their of protein coding host genes. Thus any current sequence analysis pipeline that ignores or fails to accurately detect nested and multimapped genes would most likely result in significant changes in read count distribution and ultimately in incorrect expression estimates, for large proportions of the transcriptome. Indeed, as sequencing depth increases and the capacity to simultaneously detect both coding and non-coding RNA improves, read assignment tools like CoCo will become essential for any sequencing analysis pipeline.

## Supporting information

## Acknowledgements

The authors are grateful to members of their groups for useful discussions, to Mathieu Durand from the RNomics platform of the UdeS for ddPCR analyses, and to Leandro Fequino for technical support. MSS and SAE are members of the RNA group and the Centre de recherche du Centre hospitalier universitaire de Sherbrooke (CRCHUS).

## Funding

This work was supported by a Natural Sciences and Engineering Research Council of Canada (NSERC) discovery grant [to MSS], a Canada Research Chair in RNA Biology and Cancer Genomics [to SAE] and the Fonds de Recherche du Québec – Santé (FRQS) Research Scholar Junior 2 Career Award [to MSS]. [GDF] was supported by a NSERC Masters scholarship. [VB] was supported by a Masters scholarship from the FRQS. Funding for open access charge: NSERC.

## Conflict of Interest

none declared.

