## Supplementary material for "CoCo: RNA-seq Read Assignment Correction for Nested Genes and Multimapped Reads"

**Supplementary Table 1: Proportion of nested genes for different classes of RNA**

| Biotype | Total genes | Nested genes | Proportion of nested genes |
|---|---|---|---|
| **miRNA** | 1567 | 938 | 0.60 |
| **scaRNA** | 53 | 30 | 0.57 |
| **snRNA** | 1899 | 642 | 0.34 |
| **snoRNA** | 962 | 592 | 0.62 |
| **tRNA** | 628 | 141 | 0.23 |
| **lincRNA** | 7532 | 253 | 0.034 |

**Supplementary Table 2: Proportion of host genes for different classes of nested genes**

| Biotype | Total genes | Host genes to small RNAs | Proportion of host genes |
|---|---|---|---|
| **Protein coding** | 20286 | 1430 | 0.070 |
| **lncRNA** | 14668 | 305 | 0.021 |
| **Pseudogene** | 14568 | 103 | 0.0071 |

**Supplementary Table 3: Primers used for qPCR validation.** The primer pairs used in Figure 4 are listed for each gene tested.

| Gene name | Forward Sequence | Reverse Sequence |
|---|---|---|
| RN7SK | CGGTCTTCGGTCAAGGGTATACG | AGCGCCTCATTTGGATGTGTCT |
| RN7SL2 | AGGTCGGAAACGGAGCAGGTC | CGGGGTCTCGCTATGTTGCT |
| SNORA44 | GGGCTGTGGCTGGTCATAGC | AAAGCTGAGTGGCAGCTTGCAG |
| SNORA63 | TAAGTGCTGTGTTGTCGTTCCCC | TATGAGACCAAGCGTCCCTGGC |
| SNORA64 | AGTTGCACTTGGCTTCACCCG | GCACCCCTCAAGGAAAGAGAGG |
| SNORA68 | GAATCACTGTTTCTTATAGCGGTGGTT | AAATTCACTTTGAGGGGCACGG |
| SNORD16 | AATTTGCGTCTTACTCTGTTCTCAGC | TCAGTAAGAATTTTCGTCAACCTTCTGTAC |
| SNORD32A | AACATTCACCATCTTTCGTTTGAGTCTCAC | GTCTCAGAGCGGTGCATGGG |
| SNORD88C | AGCACTGGGCTCTGATCACCC | CCTCAGACCCCCAGGTGTCAA |

### Supplementary Table 4: Parameter values used for read assignment tools considered

| Tool | Pipeline name in text | Parameter values | Reference [a] |
|---|---|---|---|
| featureCounts | featureCounts_standard | --minOverlap 10 --largestOverlap -s 1 -C -T 24 -p | Liao et al 2013 |
| featureCounts | featureCounts_optimized | --minOverlap 10 --largestOverlap -s 1 -C -T 24 -p -O -M --fraction | Liao et al 2013 |
| RSEM | RSEM | convert-sam-for-rsem -p 23 && rsem-calculate-expression --strand-specific -p 23 --alignments --paired-end | Li et al 2010 |
| HTSeq | HTSeq | -f sam -r name -m intersection-strict --stranded=yes | Anders et al 2015 |
| Cufflinks | Cufflinks | -p 24 --GTF | Trapnell et al 2012 |
| STAR | STAR | --runMode alignReads --runThreadN 46 --quantMode GeneCounts --readFilesCommand zcat --outReadsUnmapped Fastx --outFilterType BySJout --outStd Log --outSAMunmapped None --outSAMtype BAM SortedByCoordinate --outSAMprimaryFlag AllBestScore --alignIntronMax 1250000 | Dobin and Gingeras 2016 |

**[a] References**


Anders, S., Pyl, P.T. and Huber, W. (2015) HTSeq--a Python framework to work with high-throughput sequencing data, *Bioinformatics*, **31**, 166-169.

Dobin, A. and Gingeras, T.R. (2016) Optimizing RNA-Seq Mapping with STAR, *Methods Mol Biol*, **1415**, 245-262.

Li, B*., et al.* (2010) RNA-Seq gene expression estimation with read mapping uncertainty, *Bioinformatics*, **26**, 493-500.

Liao, Y., Smyth, G.K. and Shi, W. (2013) The Subread aligner: fast, accurate and scalable read mapping by seed-and-vote, *Nucleic Acids Res*, **41**, e108.

Trapnell, C*., et al.* (2012) Differential gene and transcript expression analysis of RNA-seq experiments with TopHat and Cufflinks, *Nat Protoc*, **7**, 562-578.

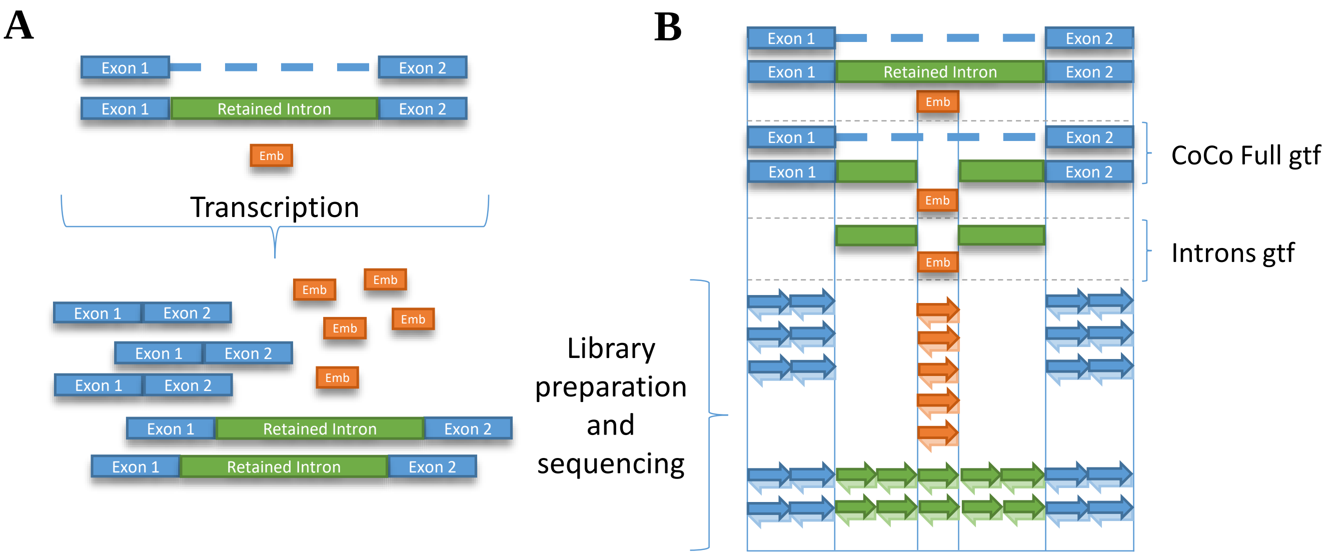


**C**

| Feature | Feature length | CoCo Full gtf | CoCo introns | Number of reassigned reads | Final counts |
|---|---|---|---|---|---|
| Embedded gene | x | 7 = 5 + 2 | 7 = 5 + 2 | - 8 * x/4x = -2 | 7 - 2 = 5 |
| Overlapping intron (non-overlapping portion) | 4x | NA | 8 | 8 * x/4x = 2 | NA |
| Host gene | y | 28 = 20 + 8 | NA | NA | 28 + 2 = 30 |

**Supplementary Figure 1 : Description of CoCo background correction for nested genes.** (A) Genomic loci encoding multiple genes (ie nested genes) can produce transcripts from different genes containing the same sequence. In the example, the sequence of transcripts from the embedded gene (orange) is identical to part of the sequence of a retained intron (green) of a host gene transcript. (B) Sequencing reads for this group of transcripts can thus come from the exons (blue arrows), retained introns (green arrows) or embedded gene transcripts (orange arrows). However, a fraction of the reads in the region in which the embedded gene overlaps the retained intron likely originate from the retained intron and not the embedded gene (stripped grey arrows). (C) To attribute the reads between the host gene and embedded gene transcripts appropriately, the context of the embedded gene is considered and used as a background to be subtracted from the total read counts attributed to the embedded gene. In the example presented, the nested gene is encoded in a retained intron. The portion of the retained intron not overlapping the embedded gene is covered by two reads over it length, and thus it is assumed that the same proportion of reads relative to its length should be assigned to the retained intron in the region overlapping the embedded gene. Thus reads are reassigned from the embedded gene to the retained intron.

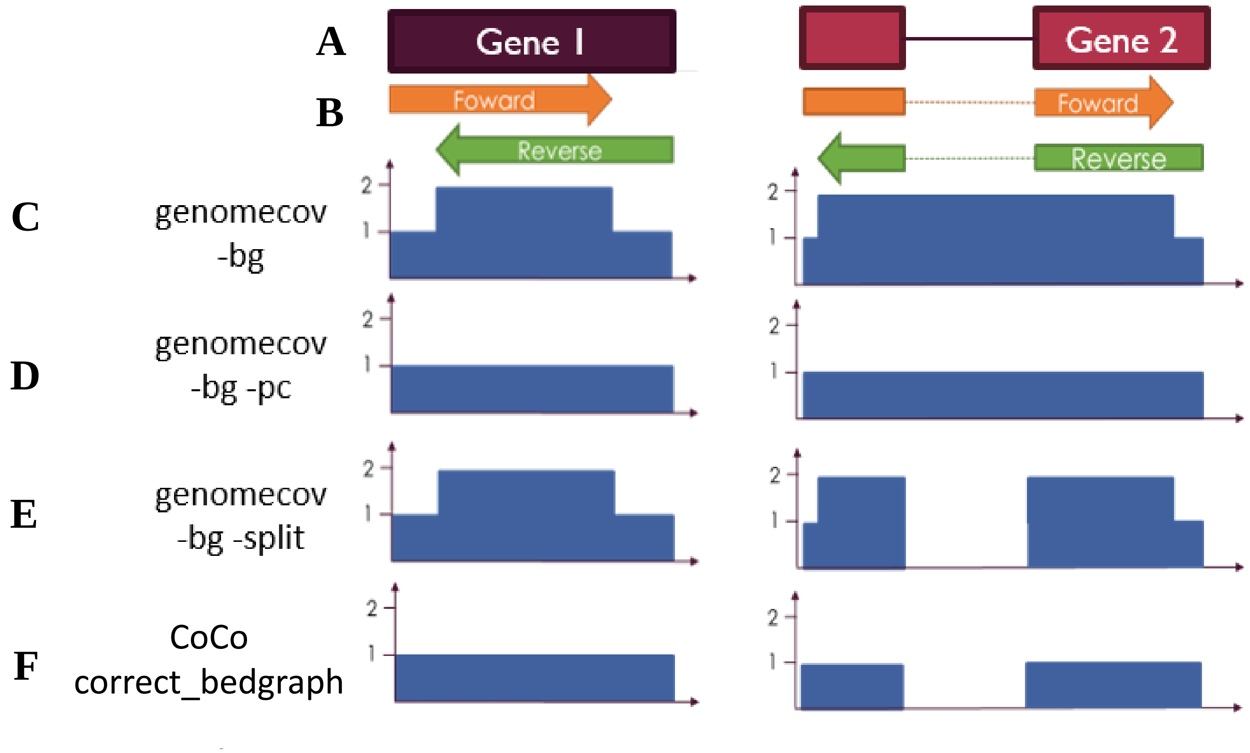


**Supplementary Figure 2 : Result of CoCo's correct_bedgraph module.** (A) Representation of two genes, having one (Gene1) and two (Gene 2) exons. (B) Representation of a read pair aligned to each gene. The read pair aligned to Gene 2 includes a splice junction. (C-F) Graphical representations of the aligned read pairs using (C-E) Bedtools genomecov with specified options and (F) CoCo's correct_bedgraph module. When a read pair doesn't contain a splice junction, the abundance is doubled where the members of the pair overlap in C and E, while the members of the pair are counted as one in D and F. When the read pair contains a splice junction, the members of the pair are individually counted in C and E. C and D report coverage in the intron, while E and F don't. Finally, in F, the pair counts as only one and the intron isn't covered.

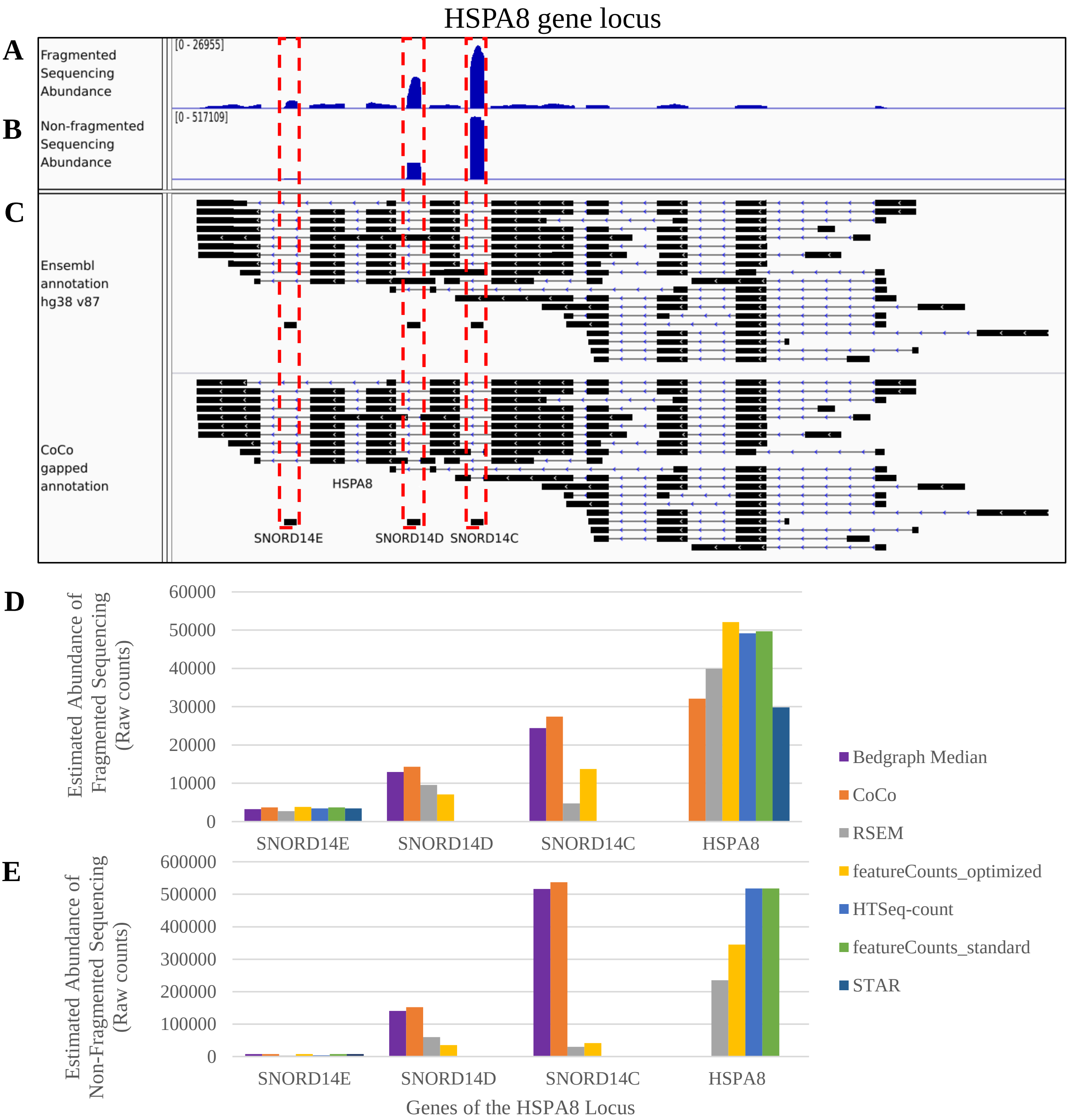


**Supplementary Figure 3**: **Example of nested C/D snoRNAs.** (A,B) Bedgraph screenshot of the HSPA8 gene locus which encodes the coding gene HSPA8 and the nested box C/D snoRNAs SNORD14E, SNORD14D and SNORD14C. The nested genes are highlighted in dashed red boxes. The upper track (panel A) is from a fragmented sequencing library while the lower track (panel B) is from a non-fragmented sequencing library. Panel C shows the annotation tracks including the original Ensembl annotation and the CoCo gapped annotation highlighting nested genes. (D, E) Raw counts obtained using the different read assignment tools considered compared to the bedgraph median values calculated, for the fragmented (D) and the non-fragmented (E) datasets. The bedgraph median is only given for the small non-coding RNAs since it is not representative of the raw counts for longer genes.

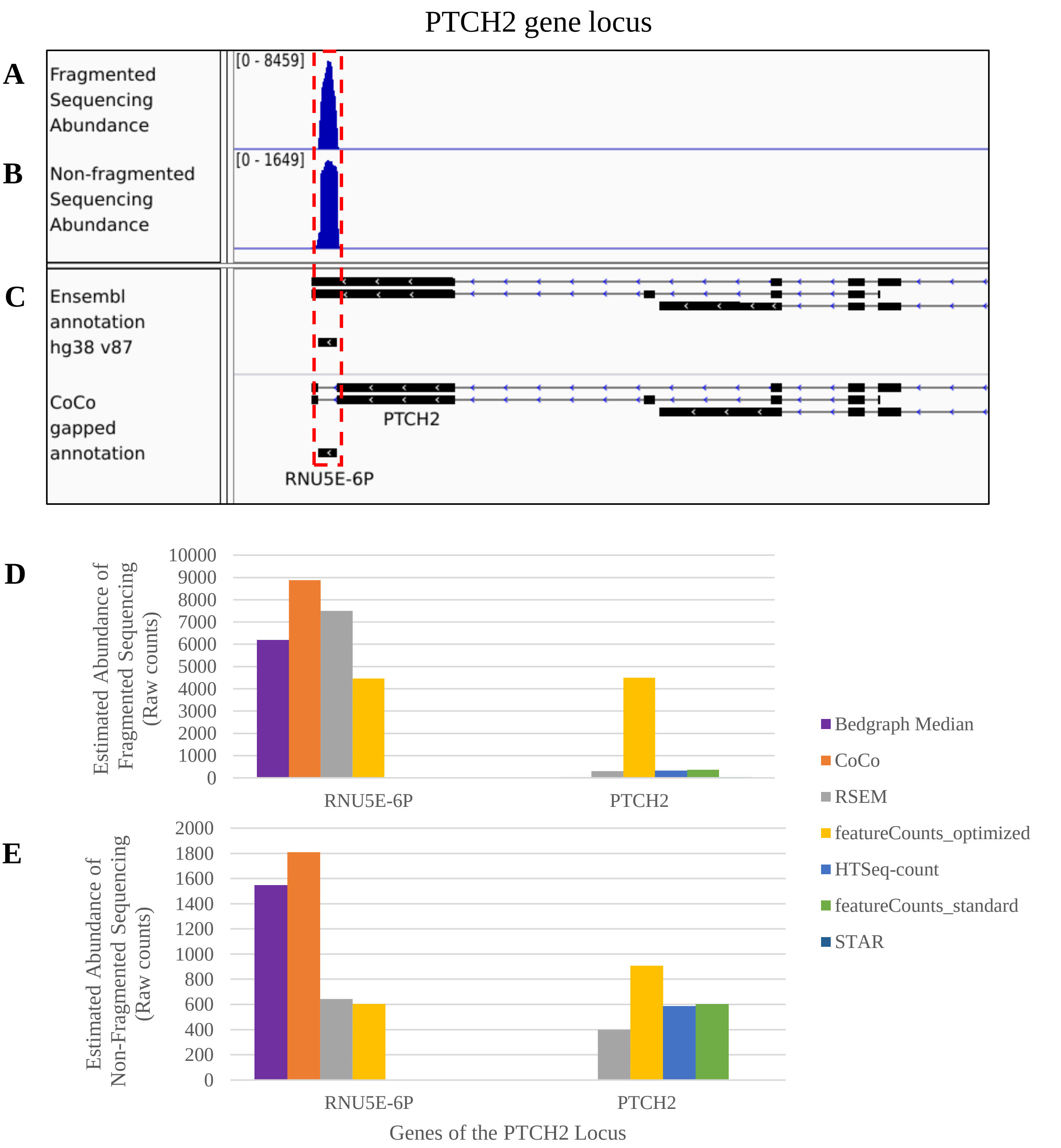


**Supplementary Figure 4 : Example of a nested snRNA.** (A,B) Bedgraph screenshot of the PTCH2 gene locus which encodes the non-coding gene PTCH2 and the nested snRNA U5 pseudogene RNU5E-6P. The nested gene is highlighted in a dashed red box. The upper track (panel A) is from a fragmented sequencing library while the lower track (panel B) is from a non-fragmented sequencing library. Panel C shows the annotation tracks including the original Ensembl annotation and the CoCo gapped annotation highlighting nested genes. (D,E) Raw counts obtained using the different read assignment tools considered compared to the bedgraph median values calculated, for the fragmented (D) and the non-fragmented (E) datasets. The bedgraph median is only given for the small non-coding RNAs since it is not representative of the raw counts for longer genes.

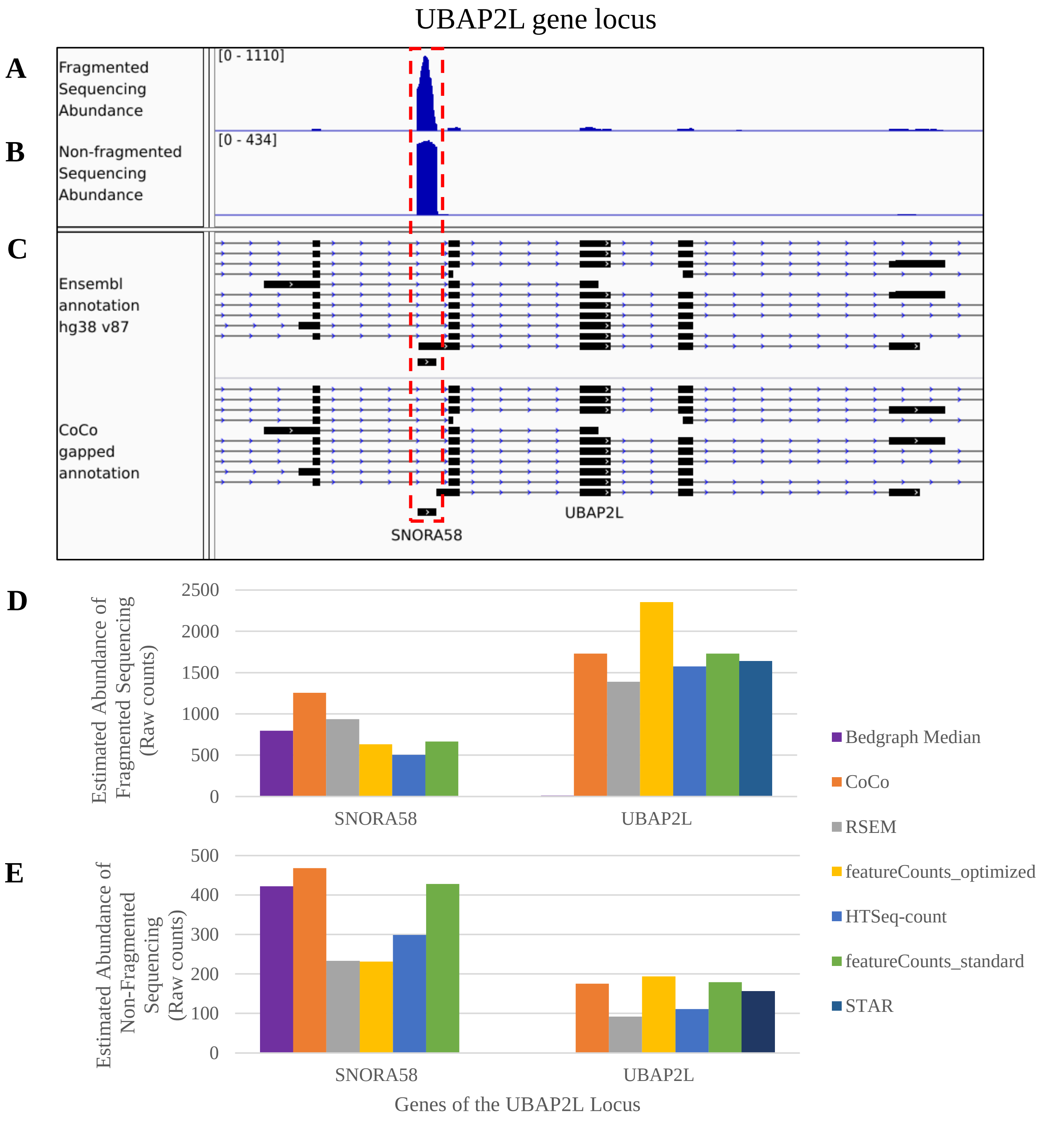


**Supplementary Figure 5: Example of a nested H/ACA snoRNA.** (A,B) Bedgraph screenshot of the UBAP2L gene locus which encodes the coding gene UBAP2L and the nested box H/ACA snoRNA SNORA58. The nested gene is highlighted in a dashed red box. The upper track (panel A) is from a fragmented sequencing library while the lower track (panel B) is from a non-fragmented sequencing library. Panel C shows the annotation tracks including the original Ensembl annotation and the CoCo gapped annotation highlighting nested genes. (D,E) Raw counts obtained using the different read assignment tools considered compared to the bedgraph median values calculated, for the fragmented (D) and the non-fragmented (E) datasets. The bedgraph median is only given for the small non-coding RNAs since it is not representative of the raw counts for longer genes.

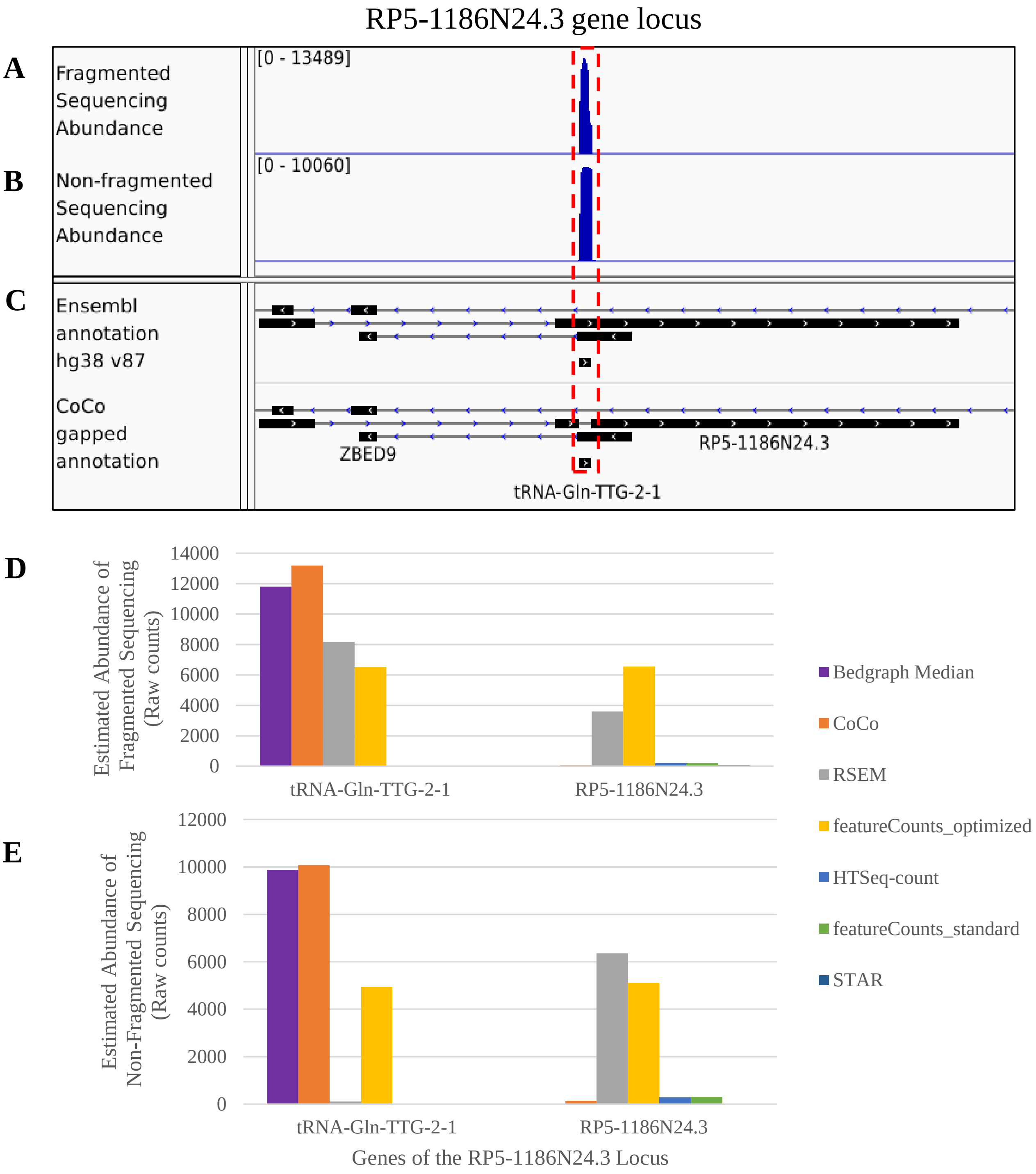


**Supplementary Figure 6: Example of a nested tRNA.** (A,B) Bedgraph screenshot of the RP5-1186N24.3 gene locus which encodes the non-coding gene RP5-1186N24.3 and the nested tRNA tRNA-Gln-TTG-2-1. The nested gene is highlighted in a dashed red box. The upper track (panel A) is from a fragmented sequencing library while the lower track (panel B) is from a non-fragmented sequencing library. Panel C shows the annotation tracks including the original Ensembl annotation and the CoCo gapped annotation highlighting nested genes. (D,E) Raw counts obtained using the different read assignment tools considered compared to the bedgraph median values calculated, for the fragmented (D) and the non-fragmented (E) datasets. The bedgraph median is only given for the small non-coding RNAs since it is not representative of the raw counts for longer genes.

SNHG21 gene locus

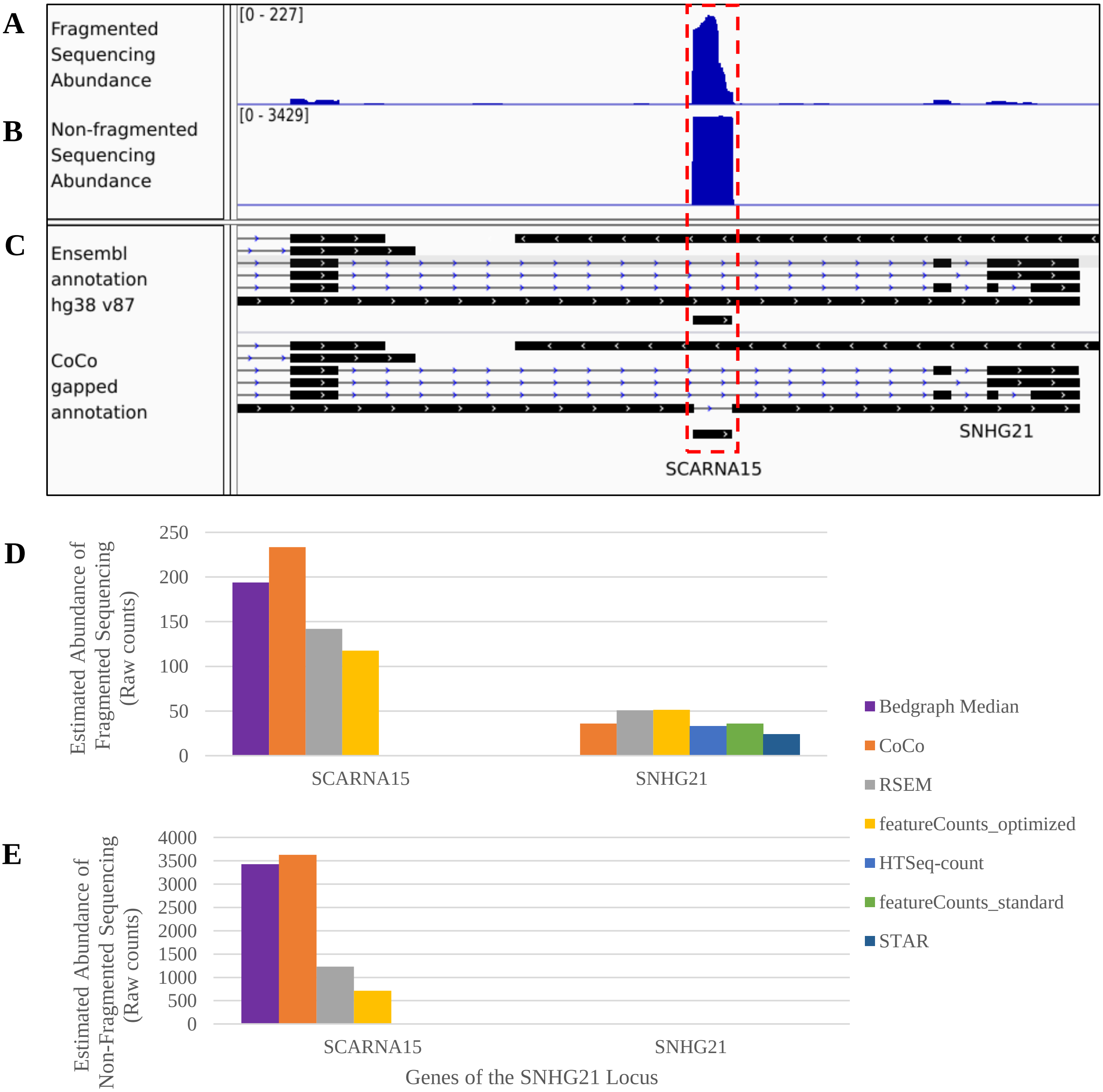


**Supplementary Figure 7: Example of a nested scaRNA.** (A,B) Bedgraph screenshot of the SNHG21 gene locus which encodes the non-coding gene SNHG21 and the nested scaRNA SCARNA15. The nested gene is highlighted in a dashed red box. The upper track (panel A) is from a fragmented sequencing library while the lower track (panel B) is from a non-fragmented sequencing library. Panel C shows the annotation tracks including the original Ensembl annotation and the CoCo gapped annotation highlighting nested genes. (D,E) Raw counts obtained using the different read assignment tools considered compared to the bedgraph median values calculated, for the fragmented (D) and the non-fragmented (E) datasets. The bedgraph median is only given for the small non-coding RNAs since it is not representative of the raw counts for longer genes.

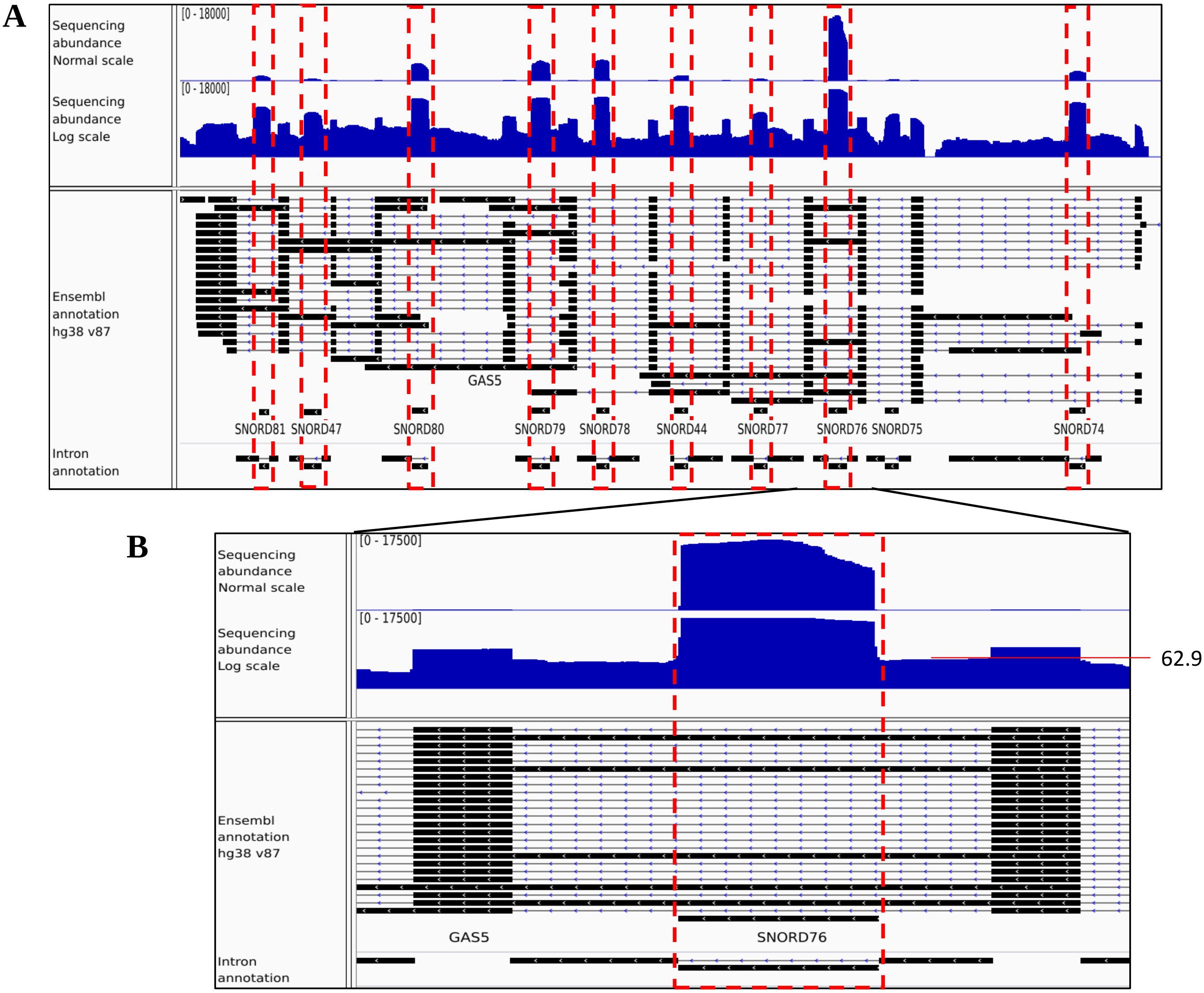


**Supplementary Figure 8: Example of the background correction for nested genes from the GAS5 locus (continued on next page).**

**C**

| Gene | Gene ID | Before Intron Correction | Number of Reassigned Reads | After Intron Correction |
|---|---|---|---|---|
| SNORD81 | NR_003938 | 1344 | -37.1 | 1306.9 |
| SNORD47 | NR_002746 | 690 | -43.8 | 646.2 |
| SNORD80 | NR_003940 | 4670 | -70.4 | 4599.6 |
| SNORD79 | NR_003939 | 5587 | -52.2 | 5534.8 |
| SNORD78 | NR_003944 | 5674 | -8.8 | 5665.3 |
| SNORD44 | NR_002750 | 1568 | -16.5 | 1551.5 |
| SNORD77 | NR_003943 | 614 | -21.2 | 592.8 |
| SNORD76 | NR_003942 | 17653 | -62.9 | 17590.1 |
| SNORD75 | NR_003941 | 523 | -16.6 | 506.4 |
| SNORD74 | NR_002579 | 2735 | -9.1 | 2725.9 |
| GAS5 | ENSG00000234741 | 2049 | 313.2 | 2362.2 |

**Supplementary Figure 8: Example of the background correction for nested genes from the GAS5 locus (continued from previous page)**. (A) Screenshot of the GAS5 locus that encodes the lncRNA GAS5 and 10 box C/D snoRNAs. The top track shows the distribution of reads for the locus in normal scale while the second track shows the distribution of reads in log scale, to make apparent the background reads of the introns. The bottom tracks show the transcript, embedded genes and intron annotations for this locus. The nested genes are highlighted in dashed red boxes. (B) Same as (A), but for a zoomed in region of the GAS5 locus, focusing on the embedded gene SNORD76 and its flanking regions. While the SNORD76 embedded gene obtains nearly a maximum of 17500, the intronic regions flanking SNORD76 obtain on average 62.9 counts as shown in the log scale track. (C) Table indicating the read counts obtained for the nested and host genes before and after background correction. The background correction removes a fraction of the reads assigned to each nested gene and reassigns them to the host gene. Because the introns are detected at a very low level, the read counts obtained after the correction are not strongly different from the read counts calculated before the correction (compare the ‘Before Intron Correction’ column to the ‘After Intron Correction’ column.

**A**

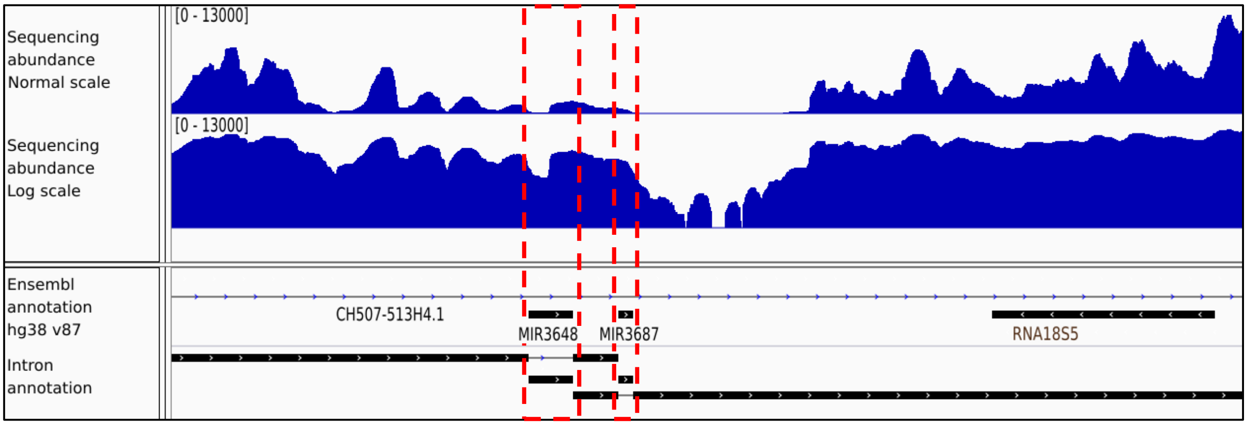


**B**

| Gene | Gene ID | Before Intron Correction | Number of Reassigned Reads | After Intron Correction |
|---|---|---|---|---|
| MIR3648 | ENSG00000275708 | 904 | -904 | 0 |
| MIR3687 | ENSG00000277437 | 587 | -587 | 0 |

**Supplementary Figure 9: Example of the background correction for nested genes from the CH507-513H4.1 locus**. (A) Screenshot of the CH507-513H4.1 locus that encodes the lncRNA CH507-513H4.1 and 2 miRNAs, miR3648 and miR3687. The top track shows the distribution of reads for the locus in normal scale while the second track shows the distribution of reads in log scale, to make apparent the background reads of the introns. The bottom tracks show the transcript, embedded genes and intron annotations for this locus. The nested genes are highlighted in dashed red boxes. (B) Table indicating the read counts obtained for the nested and host genes before and after background correction. Because the background levels are at least as high as the nested gene counts, the nested gene counts are corrected to 0 and these miRNAs are considered to be not detected in this experiment (compare the ‘Before Intron Correction’ column to the ‘After Intron Correction’ column.

**A**

```
SNORD103A   GAACTCATGAGCCCTTCTCAATTGAGGGGTTGGTTGGCCATTGTCTGGCAATGATGACCCACTTGCCCTCACTGAGAACAAAGTTC
SNORD103B   TGGGCTCTGAACTCTTCTCAATTGAGGGGTTGGTTGGCCATTGTCTGGCAATGATGACCCACTTGCCCTCACTGAGAACAAAGTTC
                   *** * *************************************************************************

SNORD103A   GGTAATGAGAATCTTTGTTAATGGACTCAAGTTCTGAGCCAGACAAGACACCCACCACCCCTGAGTCAAGCTAAACCAATCCAAA
SNORD103B   GGTAATGAGAATCTTTGTTAATGGACTCAAGTTCTGAGCCAGACAAGACACCCACCACCCCTGAGTCAAGCTAAACCAATCCAAA
            *************************************************************************************
```

**B**

| Gene | Ratio of Uniquely Mapped Reads | Number of Multimapped Read Pairs | Distribution of Multimapped Read Pairs | Total Read Pair Counts |
|---|---|---|---|---|
| SNORD103A | 50 (0.57) | 2314 | 0.57*2314 = 1330 | 50+1330 = 1380 |
| SNORD103B | 37 (0.43) | | 0.43*2314 = 984 | 37+984 = 1021 |
| Total | 87 (1) | 2314 | 2314 | 2401 |

**C**

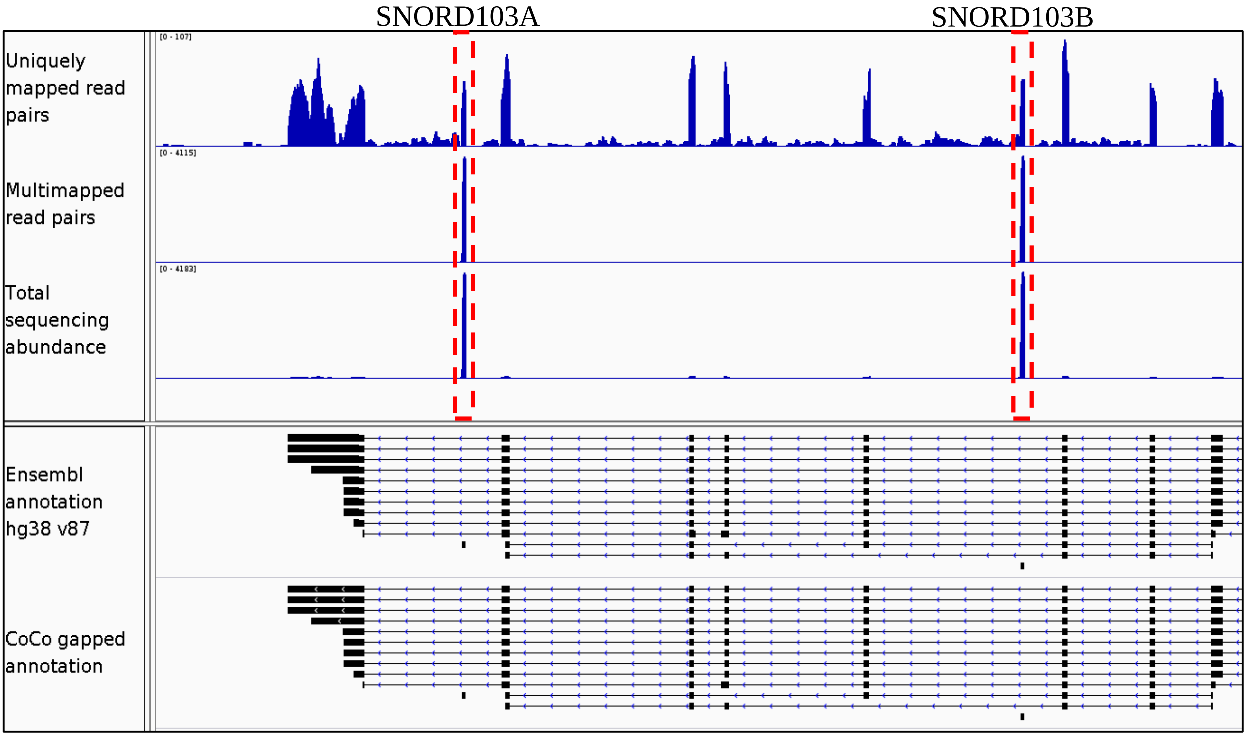


**Supplementary Figure 10 : Example of CoCo based correction for multimapped read groups.** (**A**) Comparison of the genomic sequences of the box C/D snoRNAs SNORD103A and SNORD103B. SNORD103A and SNORD103B, located in two separate introns within their host gene PUM1, have identical mature sequence (shown in black) but non-identical genomic flanking sequences (shown in red, 40 nucleotides on either side). (**B**) Example of CoCo multimapped read distribution for fragmented datasets. CoCo considers the uniquely mapped counts for SNORD103A and SNORD103B and distributes the multimapped read pairs according to the uniquely mapped ratio. (**C**) Bedgraph showing the distribution of both unique and multimapped read pairs for SNORD103A and SNORD103B. The positions of the snoRNA reads are indicated on top (highlighted in dashed red boxes) and the Ensembl and CoCo gapped annotation are shown at bottom.

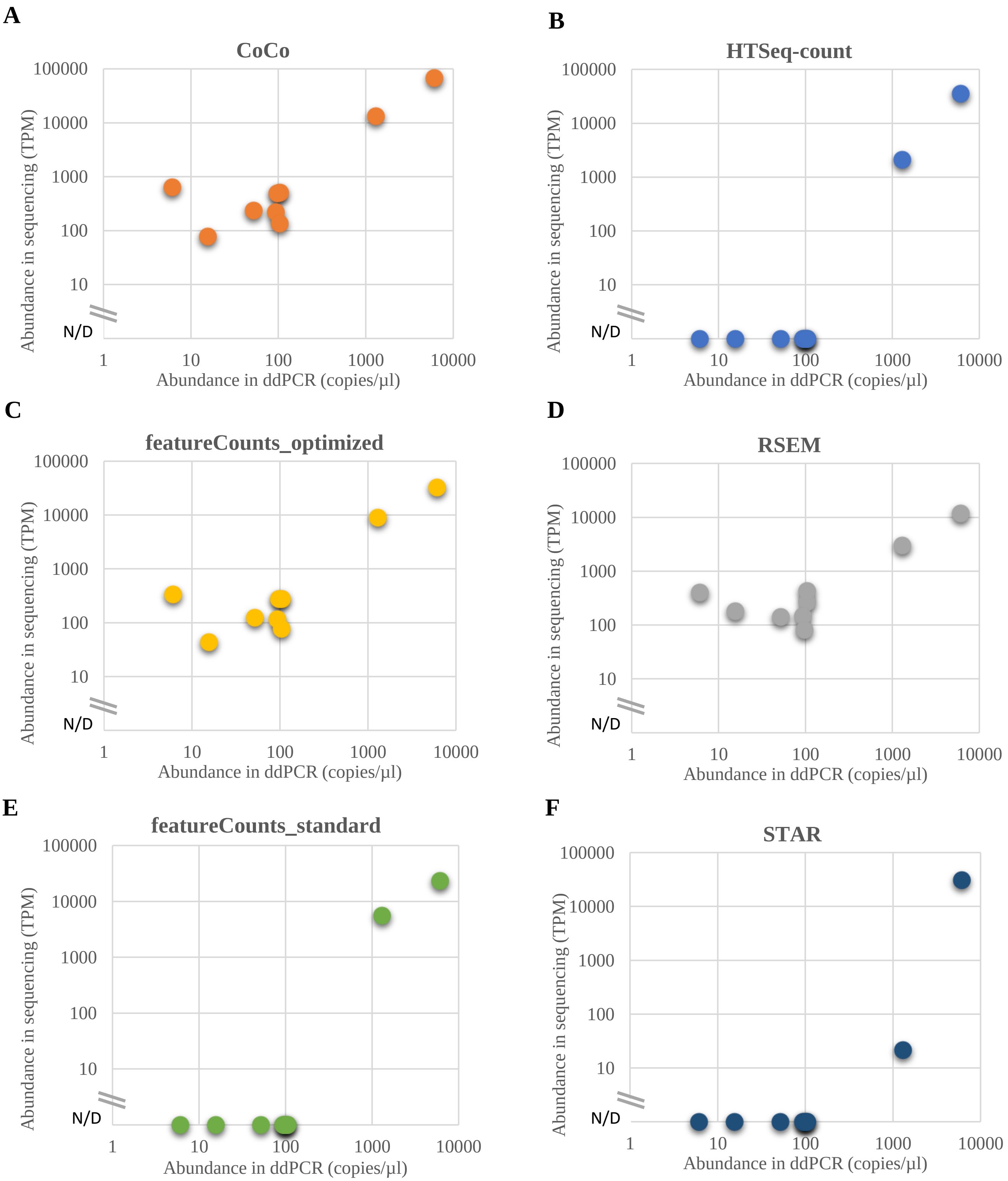


**Supplementary Figure 11: Comparison between ddPCR and different read assignment tool estimates for chosen nested and multimapped genes.** Shown are the estimated abundance of 9 nested or multimapped genes by sequencing as a function of ddPCR using the CoCo pipeline (A), HTSeq (B), featureCounts with optimized parameter values (C), RSEM (D), featureCounts with standard parameter values (E) and STAR (F). N/D in sequencing abundance stands for “Not Detected”.

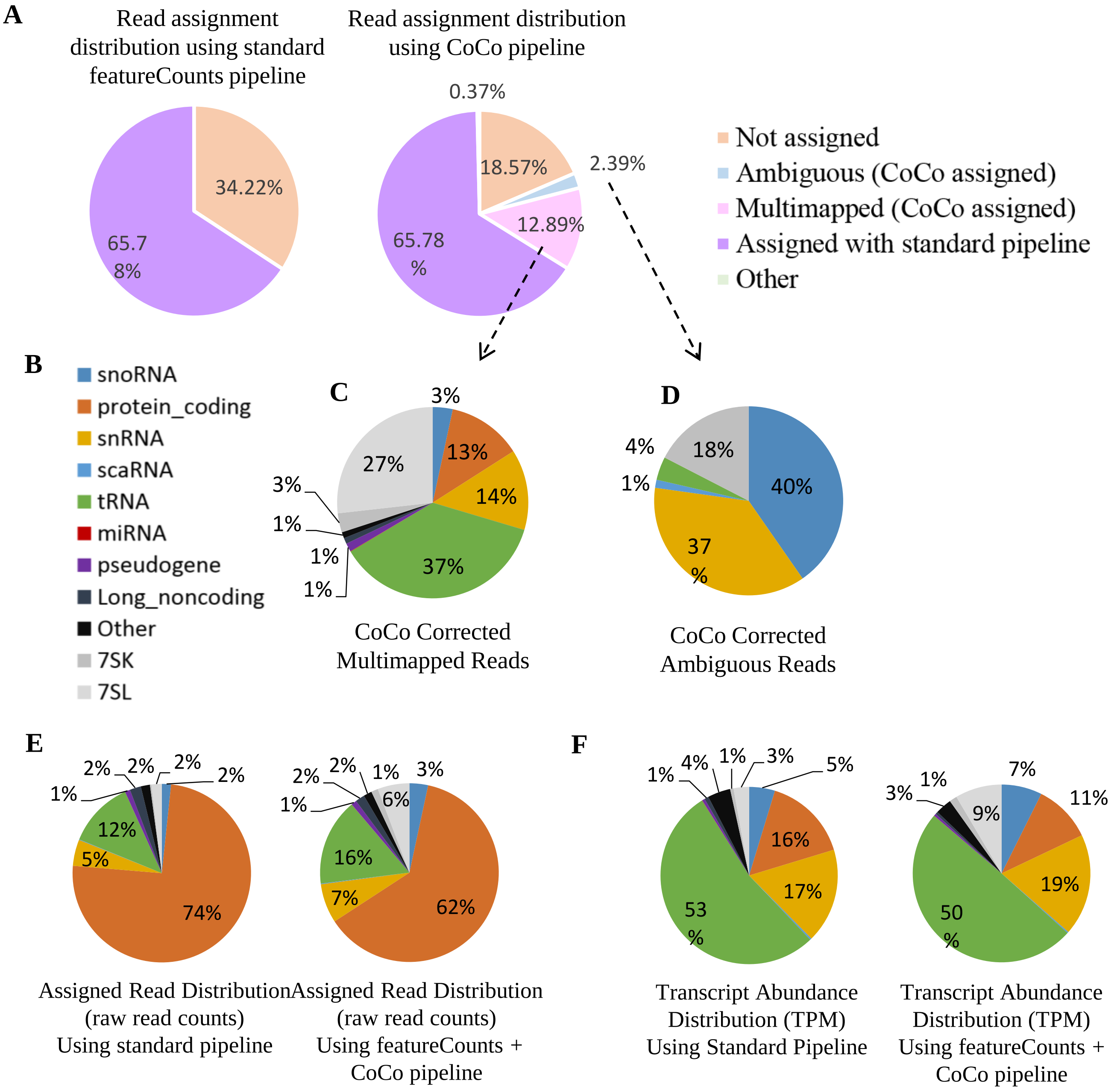


**Supplementary Figure 12: Impact of the CoCo pipeline on the distribution of read assignments and the prediction of relative RNA abundance, for fragmented datasets.** (**A**) Distribution of read assignment status as provided by the standard pipeline (left) or the CoCo pipeline (right). The color legend is shown to the right. Note that the ambiguous and multimapped reads are assigned only by CoCo. (**B**) Color legend for the pie charts in panels C-F representing the different biotypes of the genes to which read pairs were assigned. (**C**) Pie chart representing the distribution of the gene biotype to which were assigned multimapped reads by CoCo. (**D**) Pie chart representing the distribution of the gene biotype to which were assigned ambiguous reads by CoCo. (**E**) Distribution of the assigned raw read counts using the standard (left) or CoCo (right) pipelines by biotype. (**F**) Distribution of the normalized transcript estimates (TPM) assigned by the standard (left) or CoCo (right) pipelines by biotype.

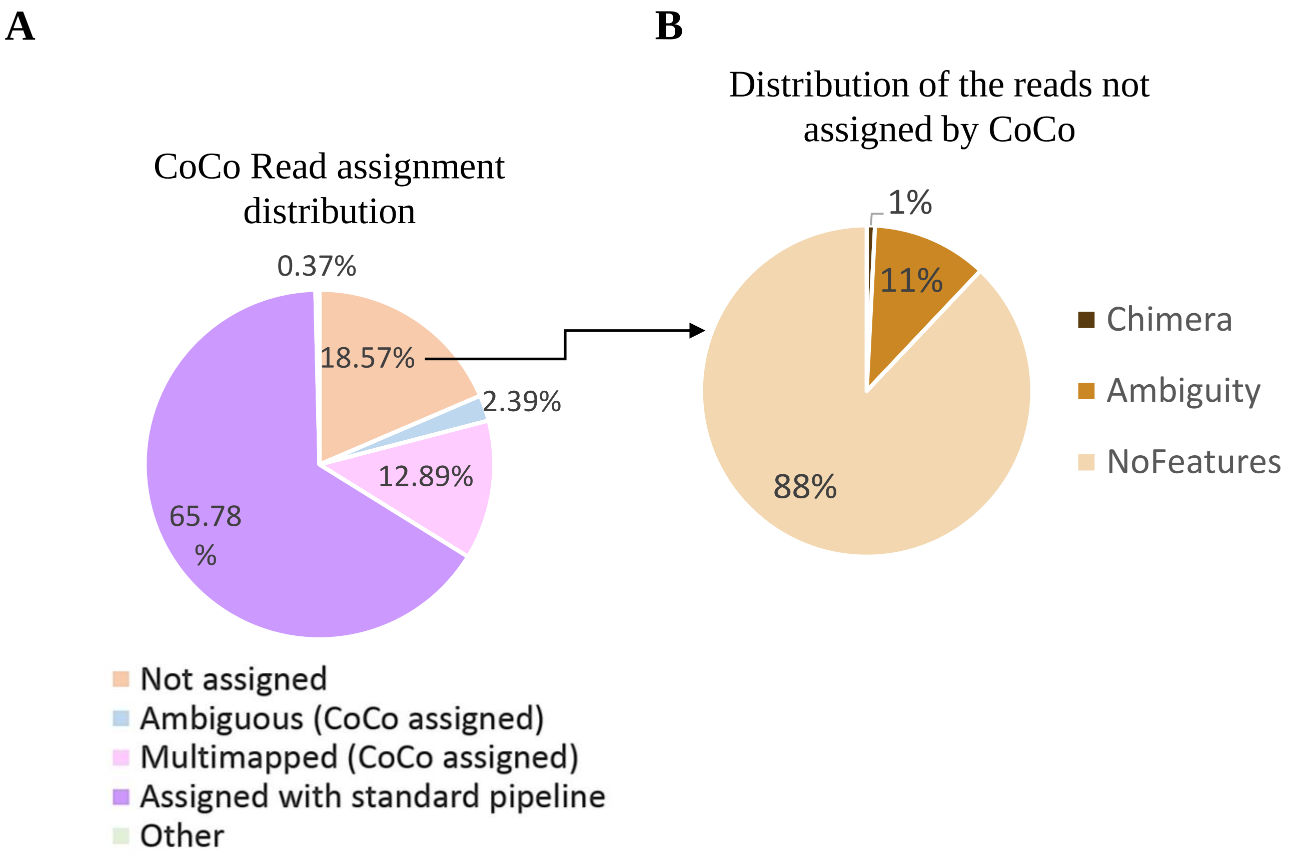


**Supplementary Figure 13: Distribution of read pairs that are not assigned by CoCo.** (**A**) Pie chart illustrating the distribution of the CoCo assigned reads. Note that 18.61% of the reads are not assigned by CoCo, (B) A pie chart showing the type of reads not assigned by CoCo. The majority of the unassigned reads (89%) align to genomic regions in which no feature is annotated. The rest of the unassigned reads are either ambiguous (aligning to multiple features) or chimeric reads.

A

B

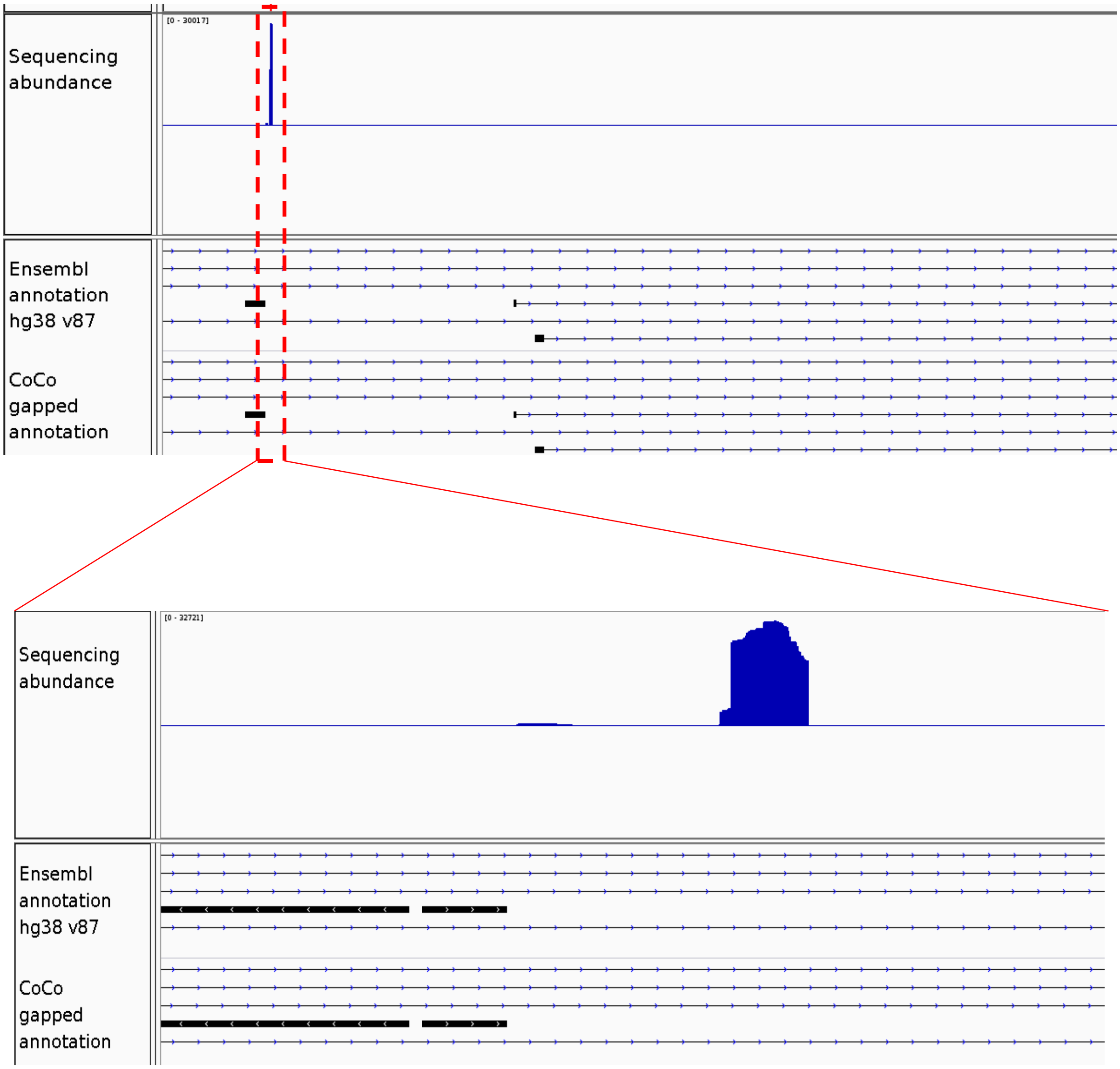


**Supplementary Figure 14: Example of reads generated from unannoted sequence.** (**A**) Shown is the bedgraph illustrating the reads accumulating in a featureless region (i.e. region with no annotated feature) within the intron of the AKAP6 gene. (**B**) Magnification of AKAP6 intronic region generating the majority of the featureless sequence reads. Note the lack of reads in the adjacent exonic sequence.

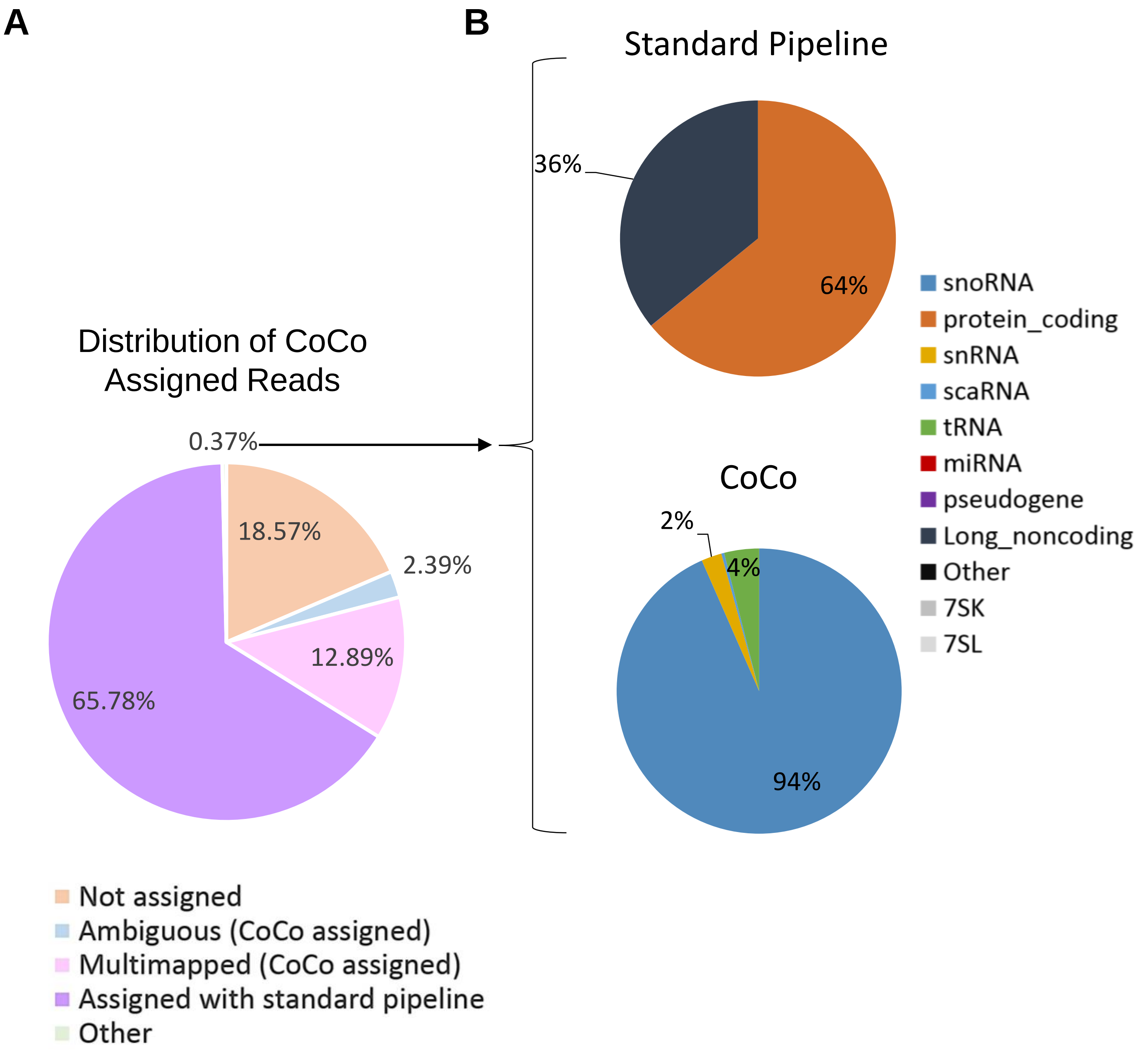


**Supplementary Figure 15: Distribution of the read pairs that are differentially assigned by CoCo.** (**A**) Coco based read distribution. The reads detected by Coco are shown in the form of a pie chart. (**B**) Distribution of the reads that are differentially assigned by CoCo and standard pipelines (featureCounts alone). The reads that are differentially assigned by the different pipelines, which represents 0.26% of the CoCo aligned read pairs or the majority of the read pairs annotated as 'Other' in panel A are shown in the form of pie charts for either the standard pipeline (upper panel) or to the CoCo pipeline (bottom panel). Note that most reads that are assigned for protein coding or lncRNA by the standard pipeline are assigned to snoRNA, tRNAs and snRNAs by CoCo.

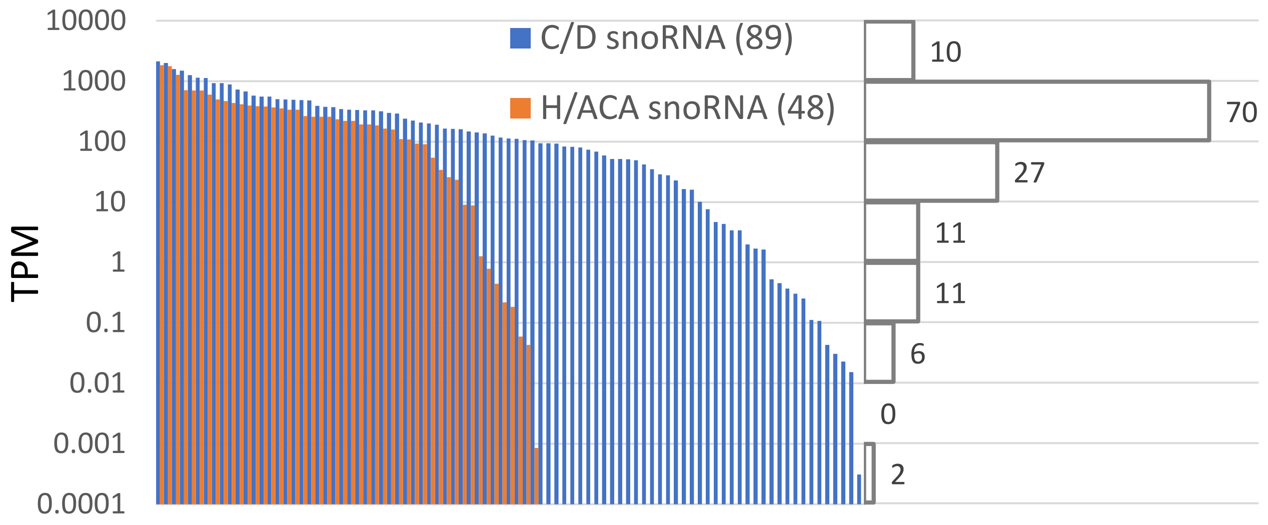


**Supplementary Figure 16: Distribution of the corrected abundance of the 130 snoRNAs detected using the CoCo pipeline but not using a standard read assignment pipeline.** The abundance for each of the 84 C/D and 46 H/ACA box snoRNA only detected by CoCo is shown as a bar chart on the left and their cumulative counts for TPM ranges is shown with bins in the horizontal histogram on the right.
